# A shape-shifting nuclease unravels structured RNA

**DOI:** 10.1101/2021.11.30.470623

**Authors:** Katarina Meze, Armend Axhemi, Dennis R. Thomas, Ahmet Doymaz, Leemor Joshua-Tor

**Author notes:** To whom correspondence should be addressed Correspondence: Leemor Joshua-Tor.

## Abstract

RNA turnover pathways ensure appropriate gene expression levels by eliminating unwanted transcripts that may otherwise interfere with cellular programs. The enzyme Dis3-like protein 2 (Dis3L2) is a 3’-5’ exoribonuclease that, through its RNA turnover activity, plays a critical role in human development^1^. Dis3L2 can independently degrade structured substrates and its targets include many coding and non-coding 3’-uridylated RNAs^1–5^. While the basis for Dis3L2’s substrate recognition has been well-characterized^6^, the mechanism of structured RNA degradation by this family of enzymes is unknown. We characterized the discrete steps of the degradation cycle by determining electron cryo-microscopy structures representing snapshots along the RNA turnover pathway and measuring kinetic parameters for single-stranded (ss) and double-stranded (ds) RNA processing. We discovered a dramatic conformational change that is triggered by the dsRNA, involving repositioning of two cold shock domains by 70 Å. This movement exposes a trihelix-linker region, which acts as a wedge to separate the two RNA strands. Furthermore, we show that the trihelix linker is critical for dsRNA, but not ssRNA, degradation. These findings reveal the conformational plasticity of this enzyme, and detail a novel mechanism of structured RNA degradation.

---

RNA quality control and turnover are vital for cellular function, yet little is known about how nucleases deal with the diverse universe of structured RNAs. Dis3L2 is an RNAse II/R family 3’-5’ hydrolytic exoribonuclease that plays an important role in development and differentiation^1,7^, cell proliferation^8–11^, calcium homeostasis^12^, and apoptosis^13,14^ by effectively removing or processing 3’ uridylated RNAs. Dis3L2 targets are oligo-uridylated by the terminal uridylyl-transferases (or TUTs)^15–17^. The specificity towards uridylated RNAs is conferred through a network of base-specific H-bonds along the protein’s extensive RNA binding surface, as demonstrated by the structure of *M. musculus* (Mm) Dis3L2 in complex with a U_13_ RNA^6^.

Genetic loss of Dis3L2 causes Perlman Syndrome, a congenital overgrowth disorder that is characterized by developmental delay, renal abnormalities, neonatal mortality, and high rates of Wilms’ tumors^1^. The first reported physiological substrates of Dis3L2 were the uridylated precursors of let-7 microRNAs (miRNA)^2,5^ which play an important role in stem-cell differentiation by silencing growth and proliferation genes such as HMGA2, MYC and Ras^18–23^. Many other non-coding RNA targets have since been reported, including other miRNAs^24,25^, tRNA fragments^17^, snRNA^26^, the intermediate of 5.8S ribosomal RNA processing 7S_B27_, the long non-coding RNA RMRP^28^, and the 7SL component of the ribonucleoprotein signal recognition particle required for ER-targeted translation^12^. The latter is most likely responsible for the Perlman syndrome phenotype, with aberrant uridylated 7SL leading to ER calcium leakage that perturbs embryonic stem cell differentiation particularly in the renal lineage^12^.

Unlike a number of structurally similar homologs, Dis3L2 can degrade structured RNAs independent of external helicase activity^1,2,4,29^. Little is known about how Dis3L2 (or other capable RNase R/II family nucleases) independently degrades structured RNA. We determined the first structures of an RNase R/II family nuclease bound to a series of structured RNA substrates and analyzed the kinetic profiles of wildtype (WT) and mutant *H. Sapiens* (Hs) Dis3L2 at single nt resolution, to reveal how this nuclease achieves highly efficient degradation of structured RNA.

## Initial Binding of Dis3L2 to Structured Substrates

To understand the pre-substrate binding state, we used cryo-electron microscopy (cryoEM) to determine the structure of RNA-free HsDis3L2 to 3.4 Å resolution (construct Dis3L2^D391N^: residues 1-858 C-terminal truncation; catalytic mutant D391N) (see Methods, Fig. 1a, b, Extended data Fig. 1a-c). RNA-free HsDis3L2 has a vase-like conformation in which three OB domains – two cold shock domains (CSDs) and an S1 domain – encircle a funnel-like tunnel that reaches into the RNA-binding domain (RNB) and leads to the active site (Fig. 1b, Extended data Fig. 1d, e). The OB domains provide a large positively charged surface, which likely acts as a landing pad for the negatively-charged RNA (Extended data Fig. 1f). This structure is very similar to the structure of the mouse Dis3L2-ssRNA complex (*M. musculus* (Mm)Dis3L2-U_13_) (RMSD 1.2 Å calculated over all Cα pairs)^6^. Thus, the apo-enzyme is preorganized to bind ssRNA.

**Figure 1.**
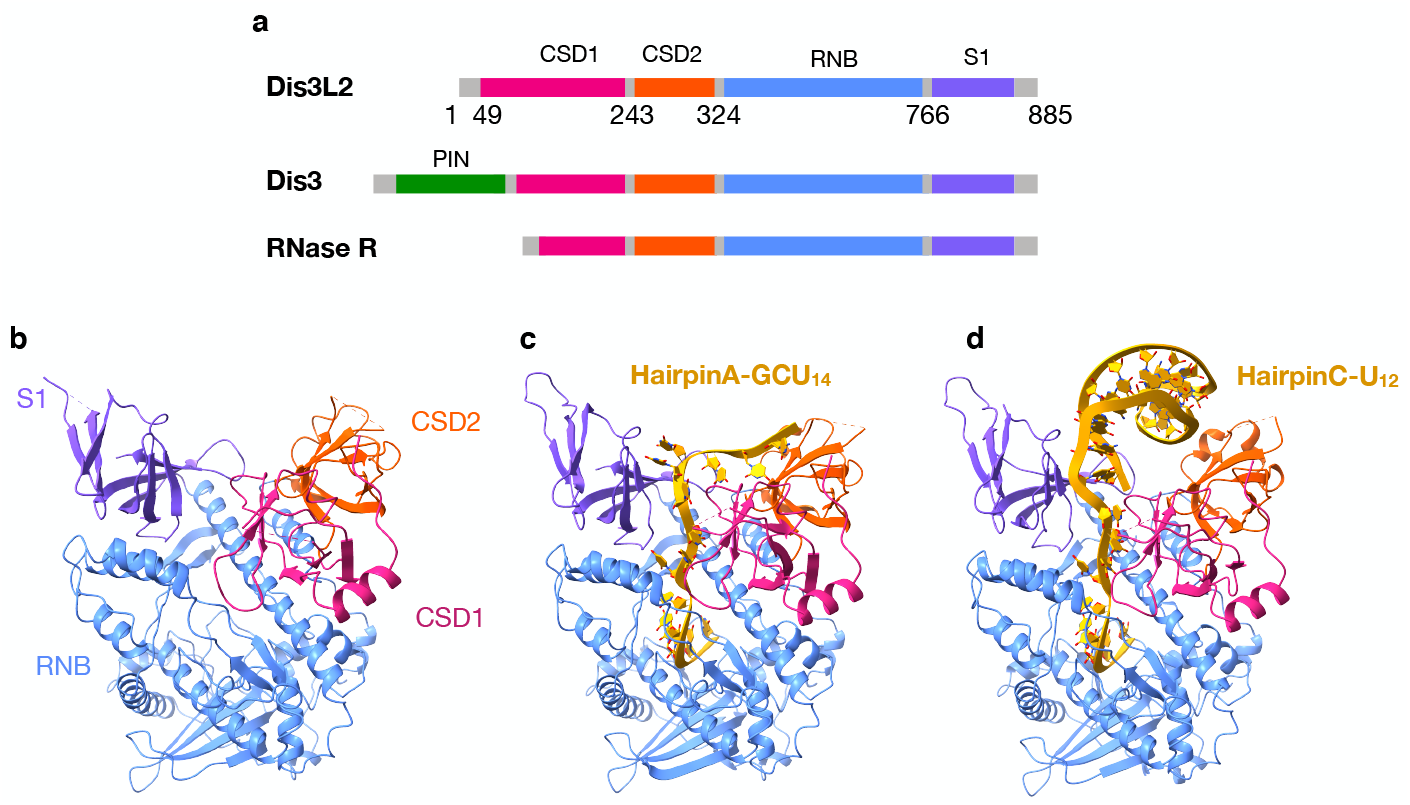
CryoEM structures of HsDis3L2 in complex with RNA substrates represent snapshots along the degradation pathway. **a**, Domain composition of Dis3L2 and homologous proteins Dis3 and RNase R: N-terminal PIN domain (green), Cold Shock Domain (CSD1-pink, CSD2-orange), RNA-binding domain (RNB-blue), S1 domain (purple). **b**, RNA-free Dis3L2^D391N^ with domain labels. **C**, Dis3L2^D391N^ in complex with hairpinA-GCU_14_. **d**, WT Dis3L2 in complex with hairpinC-U_12_.

To probe the initial binding of Dis3L2 to structured substrates we designed a short-hairpin RNA mimicking the base of the pre-let-7g stem, with a UUCG tetraloop for stability, and a 3’ -GC(U)_14_ (16nt) overhang as the uridylated tail (hairpinA-GCU_14_, Extended Data Fig. 2a). The resulting 3.1 Å cryoEM structure of the Dis3L2^D391N^-hairpinA-GCU_14_ complex revealed that Dis3L2 maintains the same vase conformation as observed in the RNA-free form (Fig. 1c, Extended data Fig. 2b-d). However, the double-helical stem of the RNA was not resolved, suggesting that dsRNA is not stably engaged by the nuclease upon initial substrate association. Nonetheless, the quality of the density allowed assignment of 15 out of the 16 nucleotides of the ss-3’ overhang. The RNA follows the same path as seen in the MmDis3L2-U_13_ structure (RMSD of 0.8 Å over C4 atom pairs), and also forms numerous base-specific hydrogen bonds with the protein (Extended data Fig. 2e-i). As in the MmDis3L2-U_13_ structure, seven nucleotides at the 3’ end are buried in the RNB domain tunnel (Extended data Fig. 2i). This tunnel can only accommodate ssRNA, which means that as Dis3L2 degrades through the ssRNA overhang and the dsRNA is brought into contact with the nuclease, the double strand will have to unwind for degradation to continue.

## 3’ tail shortening brings the RNA double helix into contact with the CSD and S1 domains

To examine the structural changes occurring upon substrate processing, we shortened the 3’ overhang to 12 uridines and further modified the stem to increase stability (hairpinC-U_12_, Extended data Fig. 3a). We obtained a 3.1 Å structure of wild-type (WT) HsDis3L2 with hairpinC-U_12_ in which the double-helical stem of the RNA hairpin is clearly visible and nestled between the two CSDs and the S1 domain (Fig. 1d, Extended data Fig. 3b-d). The basal junction of the hairpin interacts with the S1 and CSD1 domains, while the apical loop interacts with CSD2 (Extended data Fig. 3e-g). At this point, when the 3’ overhang is 12-nt long, the 5’ end, at the ss-ds junction, moves towards a loop in CSD1 (N76-H81), suggesting that further tail shortening would lead to a steric clash between the dsRNA stem and CSD1.

## RNA shortening triggers a drastic conformational rearrangement of Dis3L2 prior to dsRNA unwinding

Next, we designed a substrate with an even shorter, 7-nucleotide, 3’-overhang (hairpinD-U_7_), since this is the minimal ssRNA length needed to reach the active site from the opening to the tunnel (Extended data Fig. 2h, i, 4a). CryoEM analysis of WT HsDis3L2 in complex with hairpinD-U_7_ resulted in a 2.8 Å structure, the highest resolution Dis3L2 structure reported to date (Fig. 2a, Extended Data figure 4b-d). Strikingly, it is immediately evident that the conformation of this complex is markedly different from the vase conformation observed thus far (Extended Data figure 4b). The two CSDs moved ∼70 Å clear to the other side of the vase rim via a hinge in the linker region between CSD2 and the RNB domain (Extended Data figure 4e, SI Movie 1). This resulted in a new conformation reminiscent of a “prong” when viewed from the side (Fig. 2a). This large rearrangement is accompanied by smaller conformational changes in the S1 and RNB. The S1 domain moves such that it angles toward the double-helix where it forms new interactions with nucleotides C15 and G16 in the backbone of the double-helix, while a loop in the RNB moves by 10Å in response to the new positioning of the CSDs (Extended data Fig. 4g, h).

**Figure 2.**
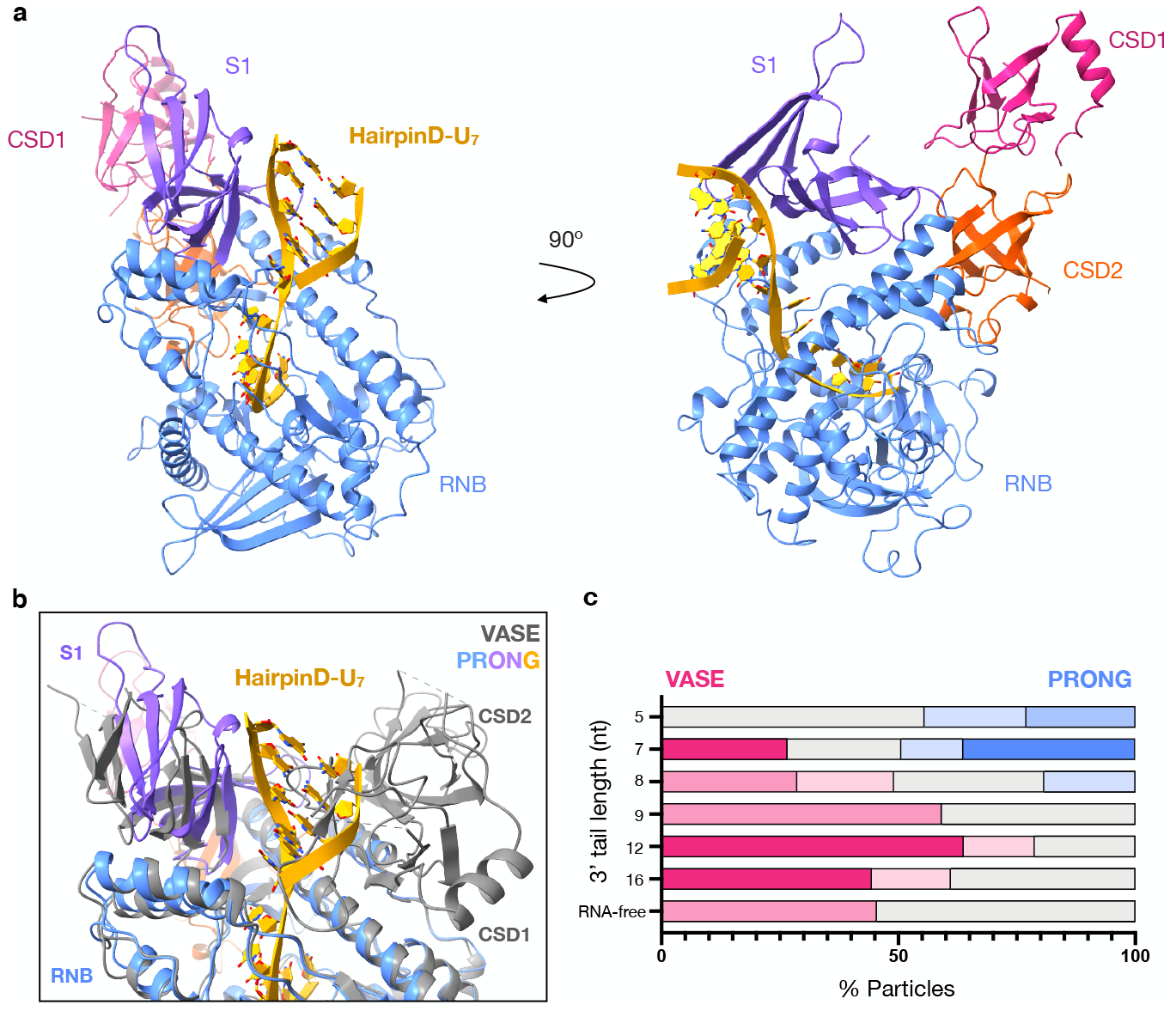
Dis3L2 undergoes a conformational change at 8 nt overhang lengths. **a**, Two views of WT Dis3L2 in complex with hairpinD-U_7_. **b**, Alignment of RNA-free Dis3L2 (gray) and Dis3L2-hairpinD-U_7_ (colored domains and RNA). The CSD domains are positioned behind S1 in the prong. **c**, Distribution of particles after heterogeneous refinement for select datasets. X-axis: % of particles in vase (pink) or prong (blue) conformation, y-axis: individual datasets RNA-free or hairpin RNA-bound Dis3L2, numbers denote the length of the 3’ overhang. The deeper color indicates higher quality 3D-reconstructions, gray indicates particles that did not contribute to a meaningful reconstruction.

The two CSDs move as a block, their relative orientations unchanged. Alignment of RNA-free and hairpinD-U_7_-Dis3L2 structures shows that the 5’ strand of the ds-RNA hairpin would clash with CSD1 (Fig 2b). Thus, it appears that upon shortening of the ss-RNA overhang, the structured portion of the RNA substrate pokes the enzyme and triggers this large rearrangement. The consequence of this movement is that it allows the structured portion of the RNA to come much closer to the RNB, while also shortening the length of the narrow tunnel to the active site by two nucleotides (Extended data Fig 5a, f). Moreover, the RNA double-helical stem is now positioned on top of the junction between a bundle of three RNB helices and a linker connecting them to the rest of the RNB domain (Fig 2a, Extended data Fig 5a, b). This junction could then act as a wedge to separate the two strands of RNA, allowing the 3’ strand to enter into the narrow tunnel. Six out of seven residues in the ss-overhang are in the same position as in the hairpinA-CGU_14_ and hairpinC-U_12_ structures, with five fully buried in the tunnel of the RNB. There is no change in the final approach to the active site. However, the 7^th^ base from the 3’ end, which is also the first ss-base, is no longer pointing towards the S1 domain and N663 as it is in the vase conformation (Extended data Fig 5c, d). Instead, it flips to stack underneath G16 of the ds-stem and forms a pseudo base-pair with R616 which emanates from the start of three α-helices in the RNB domain (Extended data Fig. 5b, e).

Upon close examination of our cryoEM data, we noticed that heterogeneous refinement yielded not only the high-resolution structure of the prong, but also a smaller 3D class representing the vase. Using a standardized analysis, we examined cryoEM data of Dis3L2 with a series of substrates of varying ss-overhang lengths and looked at their particle distributions between 3D classes after heterogeneous refinement (Methods, Extended data Fig. 6a). This analysis revealed the point in which this drastic conformational change occurs. The vase conformation is the only one observed in the RNA-free form and with long ss-3’-overhangs (Fig. 2c, Extended data Fig. 6b). The prong conformation is first observed when the overhang is 8-nt long, and is the only conformation observed when the overhang length is shortened to 5 nt (Figure 2c, Extended data Fig. 6b). This illustrates the shape-shifting nature of the enzyme to enable the degradation of structured RNA substrates, and suggests that this dramatic conformational change is triggered by the RNA when the overhang is roughly 8-nt long.

## Dis3L2 degrades structured substrates with high processivity

To quantitatively understand how the structural features described above impact Dis3L2’s function, we carried out pre-steady state kinetic assays and pulse chase experiments to measure Processivity (*P*), and elementary rate constants for RNA-binding (association: *k*_*on*_, dissociation: *k*_*off*_) and degradation (forward step: *k*_*f*_) for wildtype and various mutant Dis3L2 at single-nt resolution (Extended data Fig. 7a-g). Using a ^32^P-5’-radiolabeled 34-nt hairpin RNA with a 7-base pair stem and a 16-nt overhang (hairpinA-GCU_14_) as a substrate, we found that WT Dis3L2 is distributive for the first step (*P* = 0.23 ± 0.003), requiring approximately 3 binding events before cleaving the first nucleotide (Fig. 3a-c). This may serve as an important checkpoint before initiation of processive degradation in the second phase, where multiple nucleotides are cleaved before a dissociation event (Fig. 3c).

**Figure 3.**
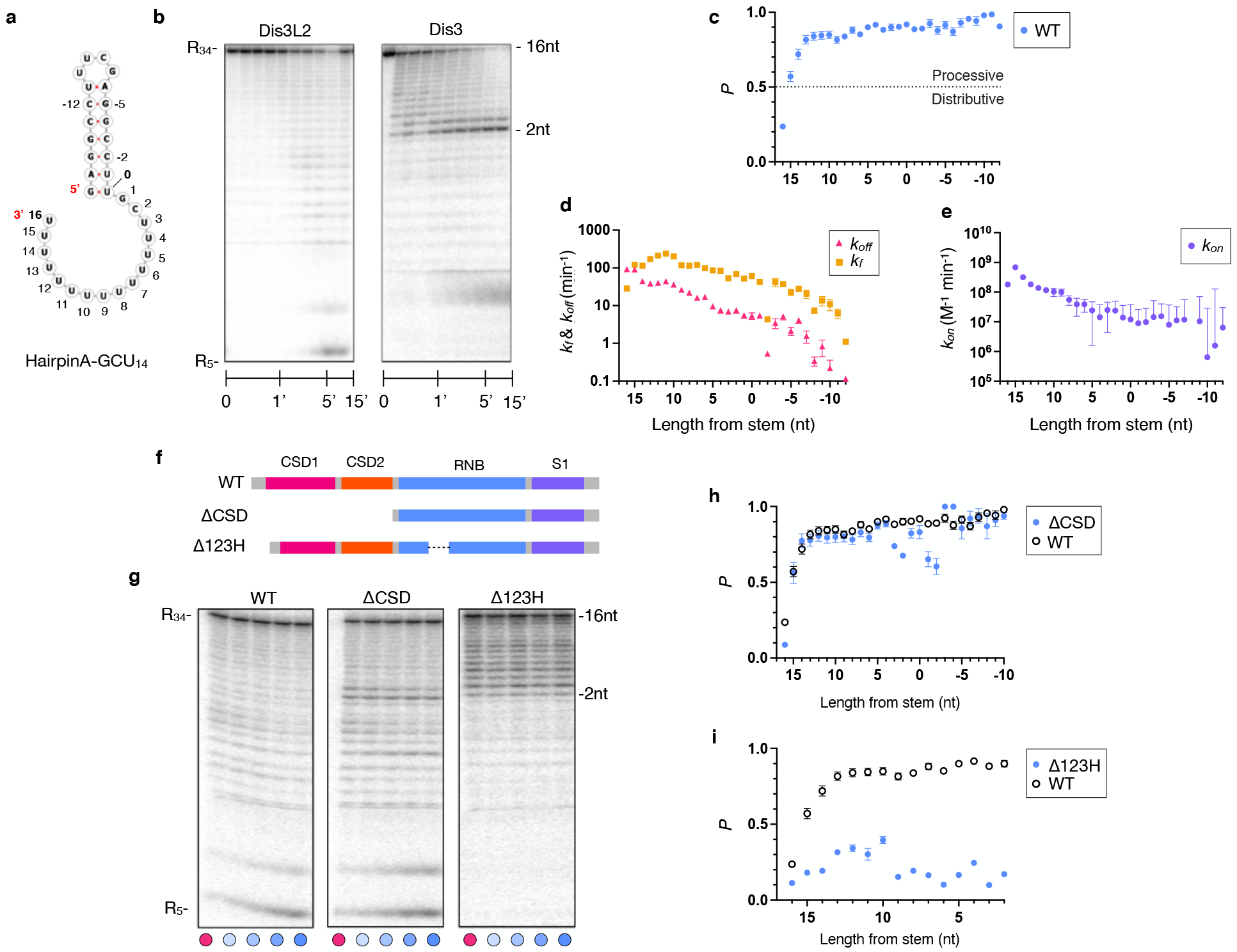
Kinetic profile of structured RNA degradation by WT and mutant HsDis3L2 at single nucleotide resolution. **a**, Schematic of hairpinA-GCU_14_, for simplicity the kinetic data is numbered from 16 (3’ end) to 0 (ss-ds junction) to denote the nucleotide position. **b**, Two representative gels from pre-steady state nuclease titration assays with 1nM 5’ P^32^ -radiolabeled hairpinA-GCU_14_ and 25nM HsDis3L2 and HsDis3, respectively. The overall length of the species and the ss-overhang length are indicated on the left and right of the panels, respectively. **c**, Processivity (P), **d**, dissociation rate constants (k_off_) and forward rate constants (*k*_*f*_), **e**, association rate constants (*k*_*on*_) of WT human Dis3L2. *k*_*on*_ datapoint x = -8 was removed due to large uncertainty value. X-axis shows number of nucleotides from the start of the double-stranded stem. **f**, Domain composition of WT human Dis3L2, ΔCSD, and Δ123H deletion mutants. **b**, Representative gels from pulse-chase reactions of WT human Dis3L2, ΔCSD, and Δ123H at 50nM concentration and 1nM radiolabeled hairpinA-GCU_14_. Cold chase was added to the reaction at the 3min timepoint to a final concentration 5000 nM. Timepoints were taken pre-chase at 3 min (pink dot), and post-chase at 4, 5, 7.5, and 10 min (blue gradient dots). Comparison of Processivity (P) for **c**, WT and ΔCSD, and **d**, WT and Δ123H. Errors for plots **d** and **e** represent standard errors of the mean (SEM), while processivity plots **c, h**, and **i** show errors calculated from the SEM of *k*_*f*_ and *k*_*off*_ (Methods).

During the course of the reaction, as the RNA is progressively shortened from the 3’ end, the *k*_*off*_ decreases (Fig. 3d). We observe a steeper decline in the *k*_*off*_ starting around 11-nt, which could reflect additional stabilizing interactions that become available when the phosphate backbone of the double-helical portion of the substrate is brought into contact with the enzyme (as was observed in the hairpinC-U_12_ and hairpinD-U_7_ structures). The *k*_on_ also decreases with RNA length, likely due to the loss of substrate interaction points (fewer 3’-U binding sites) necessary for stable association (Fig. 3e). Interestingly, the *k*_f_ varies with the stage of substrate degradation (Fig. 3d). After an initial slow step, *k*_f_ increases as the enzyme degrades through the ss-overhang. When the overhang is shortened to 11 nt, *k*_f_ peaks and then begins to fall as the enzyme encounters the dsRNA. This suggests that catalysis and/or translocation slow down as the dsRNA is engaged and later unwound, as *k*_f_ reflects the slower of these two processes. Overall, *k*_*off*_ has the dominant effect on processivity across all intermediate species (Fig. 3c).

We compared this profile to that of HsDis3, an exosome associated nuclease of this family that, in contrast to Dis3L2, cannot independently degrade structured substrates (Extended data Fig. 8a)^4^. HsDis3 appears to bind the oligo-U tailed substrate much tighter and enters directly into processive degradation (Extended data Fig. 8b, c). It maintains high processivity up until the ss-overhang reaches 3 to 2-nts in length, at which point there is a drastic decrease in processivity as a result of a large increase in the dissociation rate (*k*_off_) (Fig 3b, Extended data Fig. 8b, e). This shows that unlike HsDis3L2, HsDis3 is not able to maintain sufficient association with the substrate once it encounters the structured portion of the substrate.

## The cold shock domains play multiple roles in substrate processing

Since the CSDs appeared to be the initial recognition sites for the RNA, but then triggered to move to the other side of the protein upon RNA processing, we tested whether they contribute predominantly to initial substrate association or ssRNA degradation. Removal of the CSDs (ΔCSD: A326-S885) led to much lower processivity for the very first step, largely due to a lower forward rate constant, indicating that the CSDs play a role in augmenting the rate of catalysis for the first nt cleavage (Fig. 3f-h, Extended data Fig. 9a). During the following ssRNA degradation steps, ΔCSD has a similar processivity to WT Dis3L2 (Fig. 3h). However, there is a significant decrease in the processivity as ΔCSD approaches the structured part of the RNA, showing a marked reduction at the 3 and 2 nt ss-overhang position, as well as at -1 and -2 nt positions, which now fall within the RNA stem. This is a result of a significant increase in the dissociation rate (*k*_off_) (Fig 3h, Extended data Fig. 9b). Thus, the CSDs contribute to both initiation of RNA degradation and maintenance of substrate association during the initial unwinding steps (Supplementary discussion).

## The RNB trihelix and linker are necessary for resolving RNA structure

The Dis3L2-hairpinD-U_7_ complex structure shows that the trihelix-linker provides the final barrier before the narrow tunnel to the active site, suggesting a role in dsRNA unwinding. Deletion of the trihelix and linker (residues P612-M669: Δ123H) had a striking effect on substrate degradation and a build-up of intermediate species was observed at lengths close to the start of the double strand of hairpinA-GCU_14_ (Fig. 3f, g, i). Δ123H never reaches the processive phase, though there is a slight increase in the processivity during initial degradation of the ssRNA overhang. When the substrate shortens to 10 nt in the overhang the dissociation rate (*k*_off_) increases significantly and the forward rate (*k*_f_) plateaus leading to a dramatic drop in processivity and a buildup of species with 3, and 2 nt overhangs (Extended data Fig. 9d, e). However, no such build-up was observed in the case of a single strand U_34_ substrate, demonstrating that the trihelix-linker module is crucial for ds-, but not ss-RNA degradation (Extended data Fig. 9g-j).

## A comprehensive model for structured RNA degradation

Combining our cryoEM and kinetic data, we propose the following model: RNA degradation by Dis3L2 proceeds via a minimum of six sequential steps: 1) substrate association and quality control, 2) initial nucleotide cleavage, 3) 3’ single-strand degradation, 4) double-strand engagement, 5) dramatic domain realignment, and 6) concurrent double-strand unwinding and degradation (Fig. 4). During the first four stages, Dis3L2 is in the vase conformation, with the S1 and CSDs positioned to form a large, positively-charged surface for the oligo-U tail of the RNA (Fig 1b, Extended data Fig. 1f). While the S1 and RNB domains provide crucial binding interactions, the CSDs enable the effective initiation of degradation by contributing to the first catalytic step (Fig. 3h, Extended data Fig. 9a). This initial step is slow, and acts as a substrate checkpoint. Once cleared, the enzyme enters the highly processive phase (Fig. 3c). When the overhang is shortened to 11-12 nt, the RNA duplex engages with the enzyme, stabilized by contacts with the S1 and CSDs (Fig. 1d). At this point, the forward rate constant begins to decrease as the base of the dsRNA hairpin gets closer to the tunnel in the RNB (Fig. 3d). When the ss-3’ overhang reaches 9-8 nt, the 5’ strand of the RNA double helix runs into CSD1 and triggers the large movement of the two CSDs to the other side of the enzyme (Fig. 2a-c, SI Movie 1). In the resulting prong conformation, the S1 angles towards the tunnel and engages the backbone of the RNA double helix which now sits over the RNB trihelix-linker (Extended data Fig. 4g). The trihelix-linker module acts as a wedge between the two RNA strands to separate them and enable the 3’ strand to enter into the narrow part of the now shortened tunnel (Extended data Fig. 5a). Strand unwinding likely initiates when the overhang reaches roughly 5 nt (Supplementary discussion). Alignment of the structure of Dis3L2 in complex with hairpinD-U_7_ with known structures of RNase R/II family nucleases suggest that most would have to undergo a similar conformational change in order to allow the ds-portion of the RNA access to the trihelix-linker wedge. Biochemical studies of *E. coli* RNase R have also demonstrated the importance of the trihelix in dsRNA degradation^30^, suggesting that the mechanism proposed here could be conserved in other members of the RNase R/II family of nucleases (Supplementary discussion). Collectively, this work unveils a new molecular mechanism for efficient, regulatory degradation of structured RNAs by a vital nuclease.

**Figure 4.**
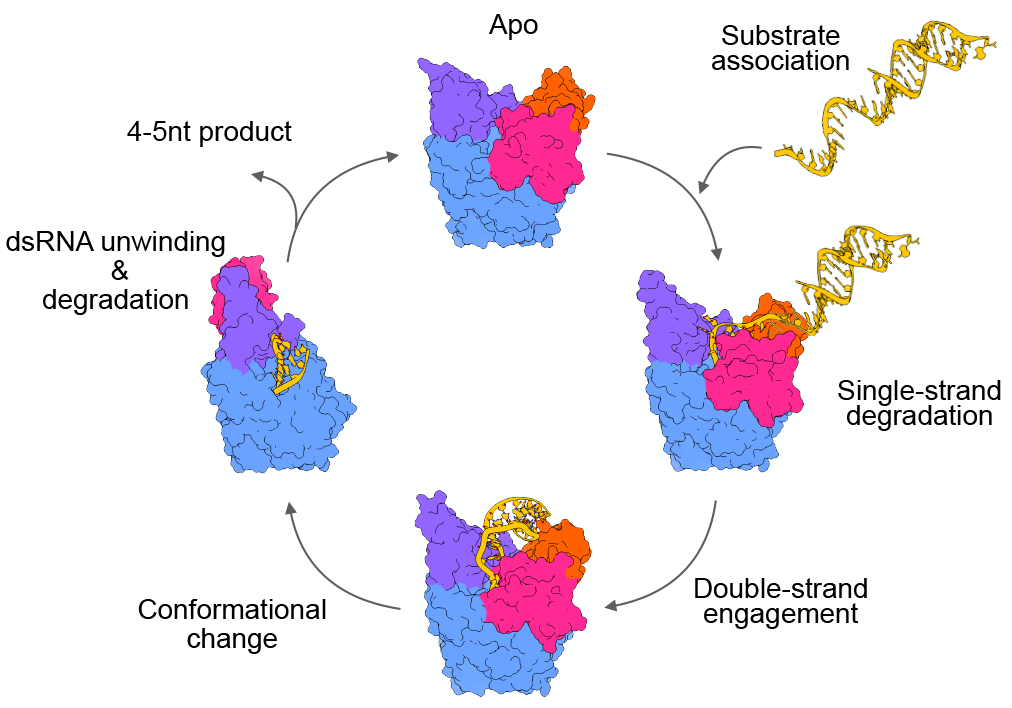
Model of structured RNA processing by Dis3L2. RNA-free Dis3L2 is pre-organized into a vase conformation to bind RNA substrates (yellow), with a 7nt deep tunnel leading to the nuclease active site. When the RNA overhang is shortened to ∼12nt, additional contacts are made to the dsRNA. Further shortening of the overhang triggers a large rearrangement of the two CSD domains (pink, orange) to the prong conformation, and allows the base of the dsRNA to access a module in the RNB domain (blue) that acts as a wedge to separate to two RNA strands and allow entry of one of the strands into the narrow tunnel leading to the active site. In this way the enzyme ensures continued RNA degradation during RNA-duplex unwinding.

## Methods

### Protein preparation

Full-length Human Dis3L2, domain deletions, point mutants, and HsDis3 were cloned as N-terminal Strep-Sumo-TEV (SST) fusion proteins in a pFL vector of the MultiBac Baculovirus expression system^31^. Benchling (https://www.benchling.com) was used for sequence analysis and primer design. Expression and purification followed a similar protocol as detailed in Faehnle *et al* 2014^6^. All constructs were expressed in SF9 cells grown in Hyclone CCM3 media (Thermo Scientific) at 27°C for 60 h. Cells were then pelleted and resuspended in wash buffer (50 mM Tris pH8, 100 mM NaCl, 5 mM DTT) and a protease inhibitor cocktail added before snap freezing with liquid N_2_ for storage at -80°C. After thawing, cells were lysed by increasing NaCl to 500 mM followed by one round of sonication. 0.1% poly-ethylene imine (PEI) was added to the lysate and cell debris cleared by 45 min of ultracentrifugation at 35,000 r.p.m. at 4°C. The solution was then incubated with Strep-Tactin superflow resin (IBA bioTAGnology) for 30 minutes while on a rolling shaker. The slurry was applied to a gravity column and washed with 20 column volumes of wash buffer before eluting the protein with 2 mM desthiobiotin in wash buffer. The SST tag was cleaved using TEV protease overnight at 4°C. Cleavage efficiency and sample purity were assessed by SDS-PAGE. The protein was then diluted to a final salt concentration of 50 mM and 25 mM HEPES pH 7.5, 5 mM DTT, and applied to a HiTrap heparin HP affinity purification column (GE Life Sciences) equilibrated in 50 mM NaCl, 25 mM HEPES pH 7.5, 5 mM DTT. The bound protein was eluted by applying a linear increasing salt gradient (0.05 M to 1 M NaCl). Pooled fractions of protein were then concentrated and loaded onto a 10/300 Superdex200 Increase gel filtration column (GE Life Sciences) equilibrated in 20 mM Hepes pH 7.5, 150 mM NaCl and 5 mM DTT. Protein purity was assessed by the quality of the chromatogram and by running SDS-PAGE gels. The concentrated sample was frozen in liquid N_2_ and stored at -80 °C.

### Cryo-EM sample and grid preparation

RNA oligos were purchased from Dharmacon and RNA secondary structure predictions were done using the Vienna RNAfold web server (http://rna.tbi.univie.ac.at) and diagrams made using Forna (Fig 3a, Extended Data Fig 2a, 3a)^32–34^. RNA hairpins were annealed by diluting into 20 mM Hepes pH 7.5, 150 mM NaCl and 5 mM DTT and heated to 95 °C for 3 minutes before stepwise cooling (95 °C, 50 °C, 30 °C, 4 °C). Complexes of human Dis3L2 and various RNAs were prepared by mixing equimolar ratios of Dis3L2 and RNA, incubating for 15 min and loading onto a 10/300 Superdex200 Increase gel filtration column (GE Life Sciences) equilibrated in the same buffer (for specific controls indicated in Extended data Fig. 6b 100 *µ*M EDTA was added to the buffer). Complex formation was evaluated by monitoring a peak shift and the ratio of absorbance at 260 and 280 nm. Fractions of the complex were then pooled and concentrated to roughly 0.5 mg/ml for Quantifoil carbon-coated Cu grids, or 0.3 mg/ml for Au-foil grids (Quantifoil, Jena, DE). 4ul of sample was applied to glow-discharged grids, and a Vitrobot plunger (Thermo Fisher Scientific) was used to freeze the grids in liquid ethane (humidity 95%, 20 °C, blot force 4, blot time 2.5s).

### Cryo-EM data acquisition and image processing

Data were collected on a 300 kV Titan Krios electron microscope at either 215,000X (0.84 Å pixel size) or 130,000X (0.64 Å pixel size) on a Gatan K2 or K3 detector equipped with an energy filter. A similar pipeline was used for all datasets (see below). CTF estimation, motion correction and particle picking were done concurrent to data collection with Warp-EM^35^. Good particles (as selected by WarpEM) were imported into CryoSPARC where 2D classification was done. A sub-selection of particles was made taking the best 2D classes with the highest resolution (below 4 Å for those that led to a high-resolution structure) and largest particle number (classes with more than 5000 particles) with protein-like features. This particle selection was then used in multi-class *ab initio* reconstruction. The classes were evaluated and then used as starting references for heterogeneous refinement, this time using all the good particles from Warp’s picking process. The best heterogeneous refinement classes and their particle subsets were then used for homogeneous refinement and in some cases non-uniform refinement in cryoSPARC (Structura Biotechnology)^36,37^. The Dis3L2-hairpinA-GCU_14_ dataset was also processed in Relion using their 3D classification, refinement, CTF refinement and particle polishing^38–40^. Two representative workflows (processing of the Dis3L2^D391N^-hairpinA-GCU_14_ and Dis3L2-hairpinD-7U dataset) are shown in SI Fig. 1.

In order to assess the distribution of particles between different Dis3L2 conformations we used the following standardized protocol for data processing: 1) particles from the datasets were picked using Warp-EM’s neural network-based picker, 2) good particles were then classified in CryoSPARC’s 2D classification, 3) the best 2D classes (as described above) were selected for *ab initio* reconstruction using 5 classes, and 4) the resulting 5 *ab initio* models were used as starting references for heterogeneous refinement using all of the good particles found by the Warp-EM picker (Extended data Fig. 5a). This allowed us to compare the proportion of particles in the full dataset that contributed to a particular Dis3L2 conformation. To ensure that the class distributions were not a result of RNA degradation, control datasets with EDTA were also analyzed and showed the same overall distribution (Extended data Fig. 5b).

### Model building and refinement

Model building and refinement were done in Coot and Phenix^41–43^. Since the mouse and human Dis3L2 proteins are extremely similar in sequence, the mouse Dis3L2 structure (pdb: 4pmw) was used as a starting reference for model building^6^. Once the reference structure was fit into the cryoEM map, Real-Space Refine (RSR) was used, using morphing, simulated annealing and rigid body fit in the first rounds^42^. After manual building and correction of geometric outliers and clashes using Coot, further rounds of refinement were done using both secondary structure restrictions as well as global minimization, refinement of atomic displacement parameters (ADP or B-factors) and local grid search. Refinements of complexes with RNA contained further base-pair and base-stacking restraints in the double-stranded regions. RNA-protein interactions were found with PDBePISA (https://www.ebi.ac.uk/pdbe/pisa/) and examined manually. Final model validation metrics are provided in Supplementary Information Table 1. Electrostatics were calculated using PyMol 2.2.3 (Schrodinger, LLC) at an ionic strength of 150 mM.

### Pre–steady-state and quasi–steady-state nuclease reactions

Nuclease reactions were performed in a temperature-controlled heat block at 20^°^C in a total volume of 40 *µ*L. Reaction mixtures containing 20 mM HEPES, pH 7.0, 50 mM NaCl, 5% glycerol, 100 *µ*M MgCl_2_, 1 mM DTT and Dis3L2 were pre-incubated for 5 minutes. Pre-steady state reactions were started by the addition of 5’-radiolabeled RNA substrate to a final concentration of 1 nM. The concentrations of Dis3L2 were in far excess of the RNA and ranged from 5 nM to 1000 nM, as indicated. Timepoints were taken at 7 s, 15 s, 30 s, 1’, 2’, 3’, 5’, 10’ and 15’, except for long time-course experiments where the times are indicated (Extended data Fig.7b, c). Reactions were quenched by the addition to an equal volume of stop buffer (80% formamide, 0.1% bromophenol blue, 0.1% xylene cyanole, 2 mM EDTA and 1.5 M urea). Samples were heated to 95 ^°^C and analyzed on sequencing gels composed of 20 % acrylamide and 7 M urea. Gels were exposed to phosphor screens overnight and scanned with a Typhoon FLA 7000 (GE Healthcare Life Sciences) imager. Bands were quantified using SAFA footprinting software and the values normalized for each lane^44^. For a typical reaction with a 34 nt substrate and 10 timepoints, we quantified all species larger than the 5 nt end-product and obtained approximately 300 datapoints for each Dis3L2 concentration.

### Pulse-chase nuclease reactions

Pulse-chase reactions were performed under conditions identical to those for pre-steady state reactions. Reactions were initiated by the addition of enzyme and allowed to proceed for a defined period of time (t_1_). At t_1_, an excess of cold scavenger RNA (x 5000-fold) was added to a final concentration of 5 *µ*M. After incubation for the indicated time (t_2_), aliquots were removed and quenched in stop buffer. Samples were analyzed on sequencing gels and processed as the above-described pre-steady state reactions.

### Calculation of kinetic parameters

Kinetic parameters were obtained using a global fit of the data from pre-steady state titrations and pulse-chase experiments. Global data fitting was performed using the Kinetic Explorer software, version 8.0 (Kintek Global)^45,46^. Initial parameters for the global fit were: observed rate constants (*k*_obs_), processivity values (*P*), *K*_*½*_ and *k*_obs_^max^ values for each reaction species. Observed rate constants (*k*_obs_) were calculated from pre-steady state experiments by fitting each experiment separately using the global data-fitting software GFIT^47^ to a model that calculates rate constants for a series of irreversible, pseudo-first-order reactions. Initial parameters for GFIT were obtained by fitting the disappearance of 34 nt substrate to a first order exponential: y = a1*exp(-b1*t) + c, where a1 is the amplitude, b1 is the observed rate constant (*k*_*obs*_) and c is the offset. Processivity values for individual degradation steps were determined from the distribution of substrate species before and after scavenger addition^48^(Extended data Fig. 7g), where Processivity (*P*) is defined as: *P* = *k*_f_ / (*k*_f_ + *k*_off_).

The equations to calculate processivity values from distributions of species are fit using a customized script using the Mathematica software package (Wolfram)^48^. To derive the *K*_*½*_ and *k*_obs_^max^ values, we fit *k*_obs_ *vs*. Dis3L2 concentration data to a binding isotherm function defined as: *k*_obs_ = (*k* ^max^ · [Dis3L2])·(*K*_*½*_^Dis3L2^ + [Dis3L2])^-1^. *K*_*½*_ is the functional equilibrium dissociation constant and *k*_obs_^max^ is the maximal observed rate constant at enzyme saturation (Extended data Fig. 7d). The data were then evaluated by plotting in GraphPad Prism version 9.1.2. (GraphPad Software). These initial parameters were used as guides in setting up a range of starting values for the elementary rate constants in a global fit to the minimal kinetic model as shown in Extended Data Fig. 7a. The *K*_½_ values were used to constrain the ratio of dissociation and association rate constants for productive binding by linking the two values as initial parameters. The *k*_obs_^max^ values were used to set boundaries on the forward rate constant, *k*_*f*_. Finally, the experimentally determined processivity values (*P*) were used as initial constraints on the ratio of *k*_*f*_ and *k*_*off*_. The global data fit was done in an iterative manner by alternating combinations of fixed and floating variables while tracking the overall χ^2^ value. The goodness of the fit, R^2^ was 0.94, calculated by plotting the experimental datasets *vs* the corresponding simulated data from the kinetic model (Extended data Fig. 7f). As an additional measure of the overall quality of fit we performed Fitspace analysis^46^ to determine the lower and upper boundaries of each kinetic parameter (Supplementary Information Table 2-4). For a typical substrate, roughly 2500 individual datapoints from enzyme titrations and 750 datapoints from pulse-chase experiments were used to calculate the 120 kinetic parameters that describe degradation of a 34 nt substrate down to 5 nt.

Errors for elementary rate constants represent standard errors of the mean from the global data fitting. Errors for compound rate constants such as processivity and K_1/2_ were calculated via error propagation formulas shown below.

**Equation 1. Error propagation formula for K_*1/2*_ and Processivity.**

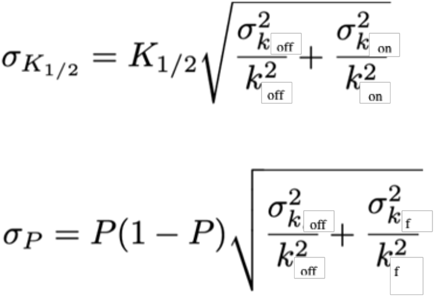

## Data availability

CryoEM data and structure coordinates will be deposited prior to publication in the EM-DB and PDB respectively.

## Acknowledgements

We thank Daniel Herschlag for detailed discussions, comments and recommendations, Justin Kinney for advice, Seraya Jones and John Scheuring for technical support, Jonathan Ipsaro for comments on the manuscript, Matt Jaremko for technical advice and other members of the Joshua-Tor lab for helpful suggestions. Cryo-EM was performed at the CSHL cryo-EM facility. We also thank the CSHL Mass Spectrometry Shared Resource, which is supported by the Cancer Center Support Grant 5P30CA045508. This work was supported by NIH grant R01-GM114147 (to L.J.), the CSHL School of Biological Sciences (to K.M., A.D. and L.J.). K.M. was supported by the Leslie C. Quick, Jr. Fellowship. L.J. is an Investigator of the Howard Hughes Medical Institute.

## Competing interest declaration

The authors have no competing interest

## Contributions

K.M. and L.J. designed the study. K.M. purified the proteins used in this work, and preformed all cryoEM sample preparation, data collection, data processing and model building. D.R.T. assisted in cryoEM data collection and A.D. contributed to the purification and cryoEM analysis of RNA-free and hairpinA-GCU_14_ HsDis3L2. A.A. designed and lead the kinetic analysis of HsDis3L2. A.A and K.M. carried out the kinetic assays and data analysis. K.M., A.A. and L.J prepared the manuscript.

## Additional information

Additional Supplementary Information and Movie are provided in separate files.

## Notes

### Competing Interest Statement

The authors have declared no competing interest.

