## Supplementary material for "A shape-shifting nuclease unravels structured RNA": Supplmentary file

*To whom correspondence should be addressed

**Extended data Figures** 3

**Supplementary Figure** 18
**Supplementary Tables** 20
**Supplementary Discussion** 22

**Supplementary Movie 1. Structural changes during the different stages of structured RNA degradation by Dis3L2.** The RNA-free Dis3L2 and domain composition in the vase conformation are shown, followed by the complex of Dis3L2 with hairpinA-GCU$_{14}$. The latter represents initial substrate binding at which point the dsRNA is not engaged by the enzyme. After shortening of the overhang to roughly 12 nt, the double-helix is cradled between the S1 and CSD domains, shown here by the structure of the complex of Dis3L2 with hairpin-U$_{12}$. When the overhang length is roughly 8 nt long, the structured portion of the RNA substrate would clash with the enzyme and therefore triggers a dramatic 70 Å domain movement of CSDs to the other side of the enzyme. A morph created in ChimeraX shows the transition between the vase and prong conformation, with the side view of the transition highlighting the large distance that the CSDs travel. This conformational change exposes the trihelix-linker that acts as a wedge to separate the two RNA strand and shortens the approach to the active site. The double-helix is further stabilized by the S1 domain in the structure of the complex of Dis3L2 with hairpinD-U$_{7}$.

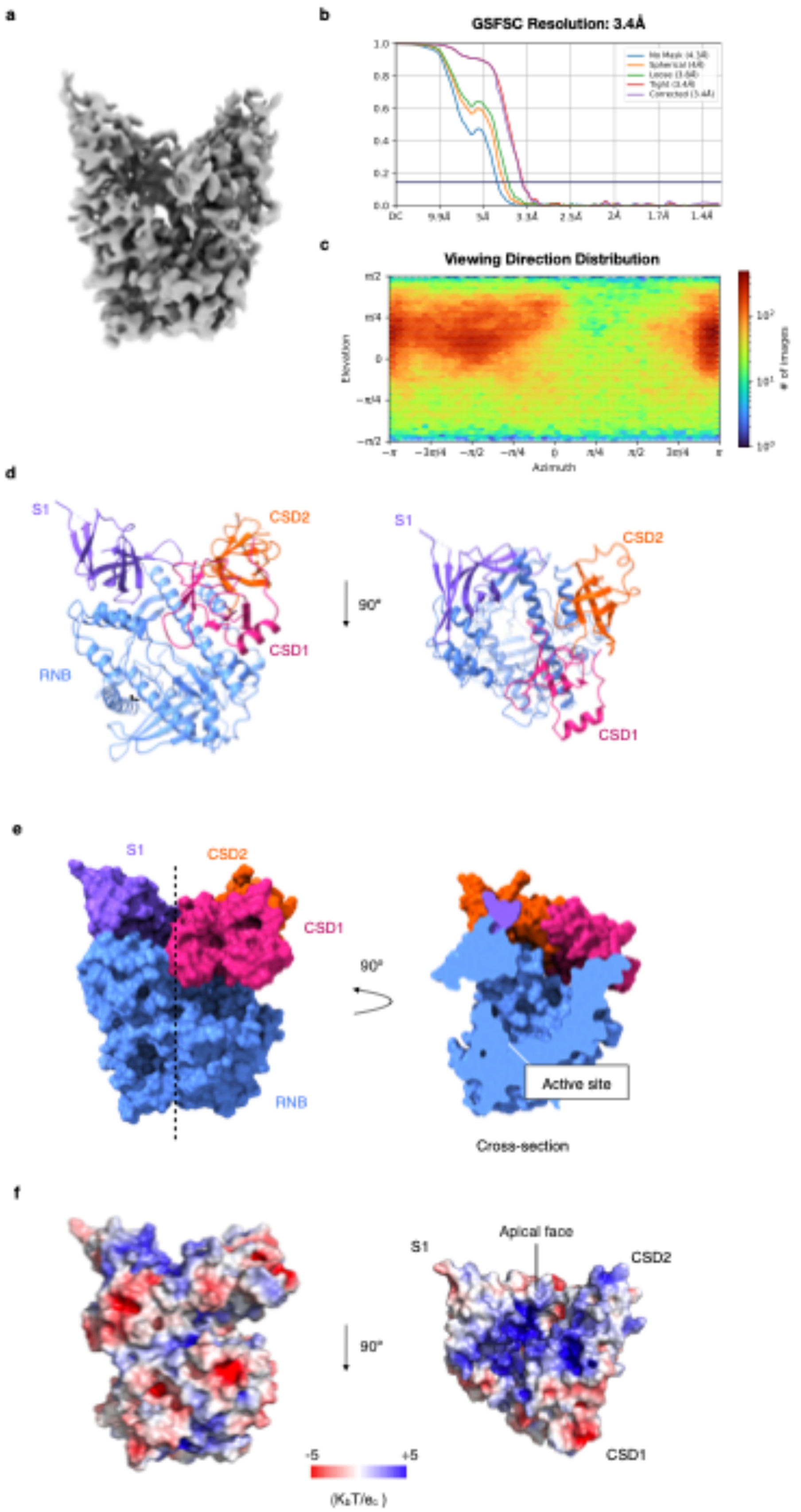
a
b
GSFSC Resolution: 3.4Å
c
Viewing Direction Distribution
Elevation
Azimuth
### of images
d
S1
CSD2
90°
CSD1
RNB
e
S1
CSD2
CSD1
90°
RNB
Active site
Cross-section
f
S1
Apical face
CSD2
90°
CSD1
-5
+5

**Extended data Fig. 1 Cryo-EM structure of RNA-free human Dis3L2**. **a**, Final 3D map of human Dis3L2 (left) with **b**, Gold Standard (0.143) Fourier Shell Correlation (GSFSC) and **c**, viewing direction distribution (bottom right). The 3D map was produced by homogeneous refinement of roughly 163-thousand particles in CryoSPARC. **d**, Structure of RNA-free Dis3L2 with domain labels and view of top or apical face (90° turn). **e**, Surface representation and cross-section (right) showing tunnel and active site. **f**, charge distribution of Dis3L2 surface and view of apical face, calculated using PyMol APBS at an ionic strength of 150 mM (Methods).

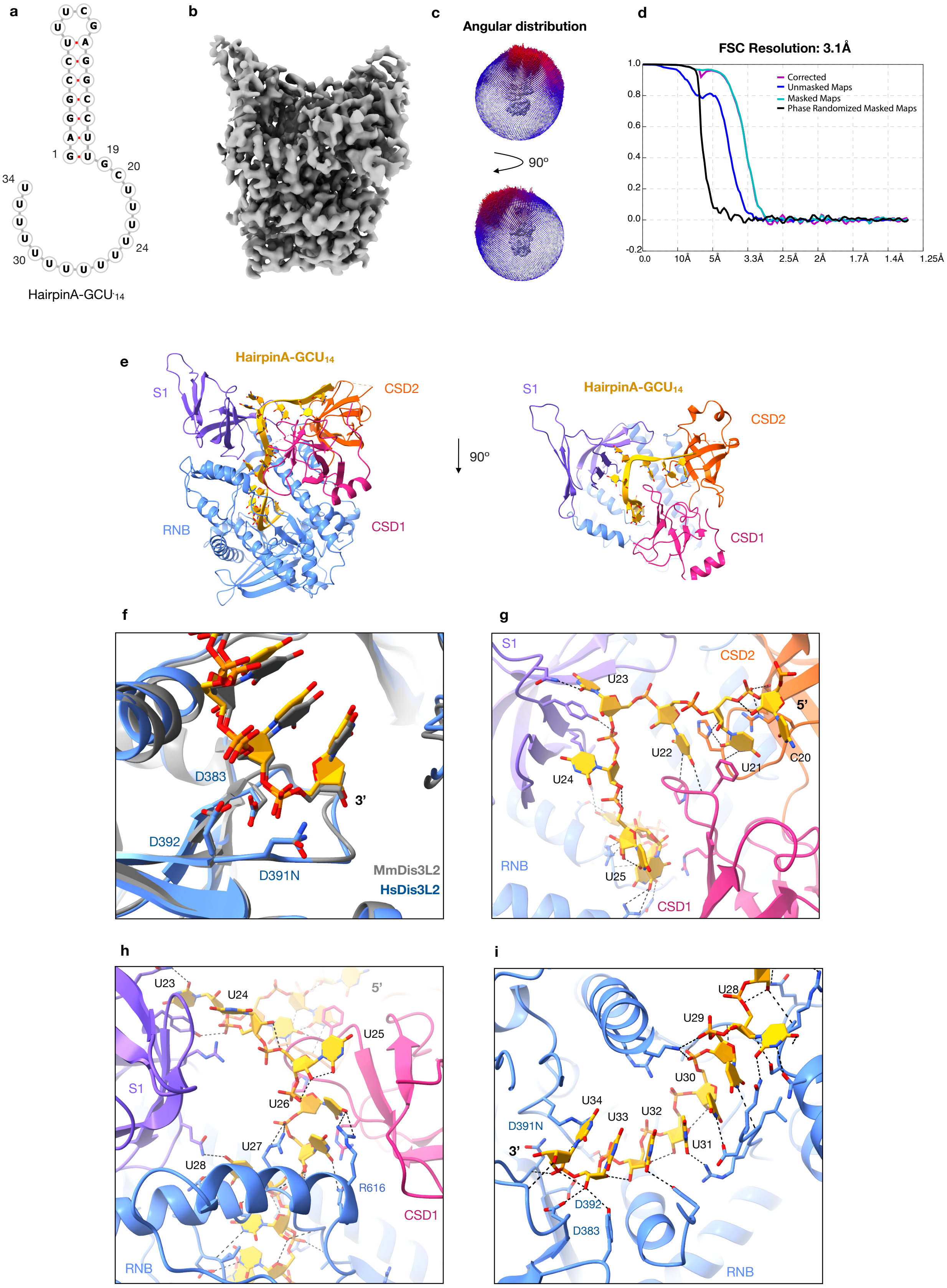
a
b
c
d
HairpinA-GCU14
Angular distribution
90°
FSC Resolution: 3.1Å
Corrected
Unmasked Maps
Masked Maps
Phase Randomized Masked Maps
e
HairpinA-GCU14
S1
CSD2
RNB
CSD1
90°
f
D383
D392
D391N
3'
MmDis3L2
HsDis3L2
g
S1
CSD2
U23
5'
U22
U21
C20
U24
RNB
U25
CSD1
h
U23
U24
5'
U25
S1
U26
U27
U28
R616
CSD1
RNB
i
U28
U29
U30
U34
U33
U32
U31
D391N
3'
D392
D383
RNB

**Extended data Fig. 2 Initial substrate binding by human Dis3L2. a**, Predicted RNA-fold for HairpinA-GCU$_{14}$. **b**, Final 3D map of human Dis3L2 in complex with hairpinA-GCU$_{14}$ with **c,** particle angular distribution and **d,** GSFSC. The final 3D map was produced by 3D refinement of roughly 101,000 particles in Relion. **e**, Structure of Dis3L2 in complex with hairpinA-GCU$_{14}$ with domains indicated, and view of top/apical face (90° turn). **f**, Alignment showing the active site of the MmDis3L2 in complex with U$_{13}$ (gray)(4pmw) and HsDis3L2 in complex with hairpinA-GCU$_{14}$. **g**, Hydrogen-bond network from C20-U25 traverses the apical face of the protein and involves both CSDs and the S1 domains. **h,** After U25, the RNA enters into the narrow portion of the channel and the H-bond network continues within the RNB domain, **i,** all the way to the active site. For C20-U32 these include base specific H-bond interactions.

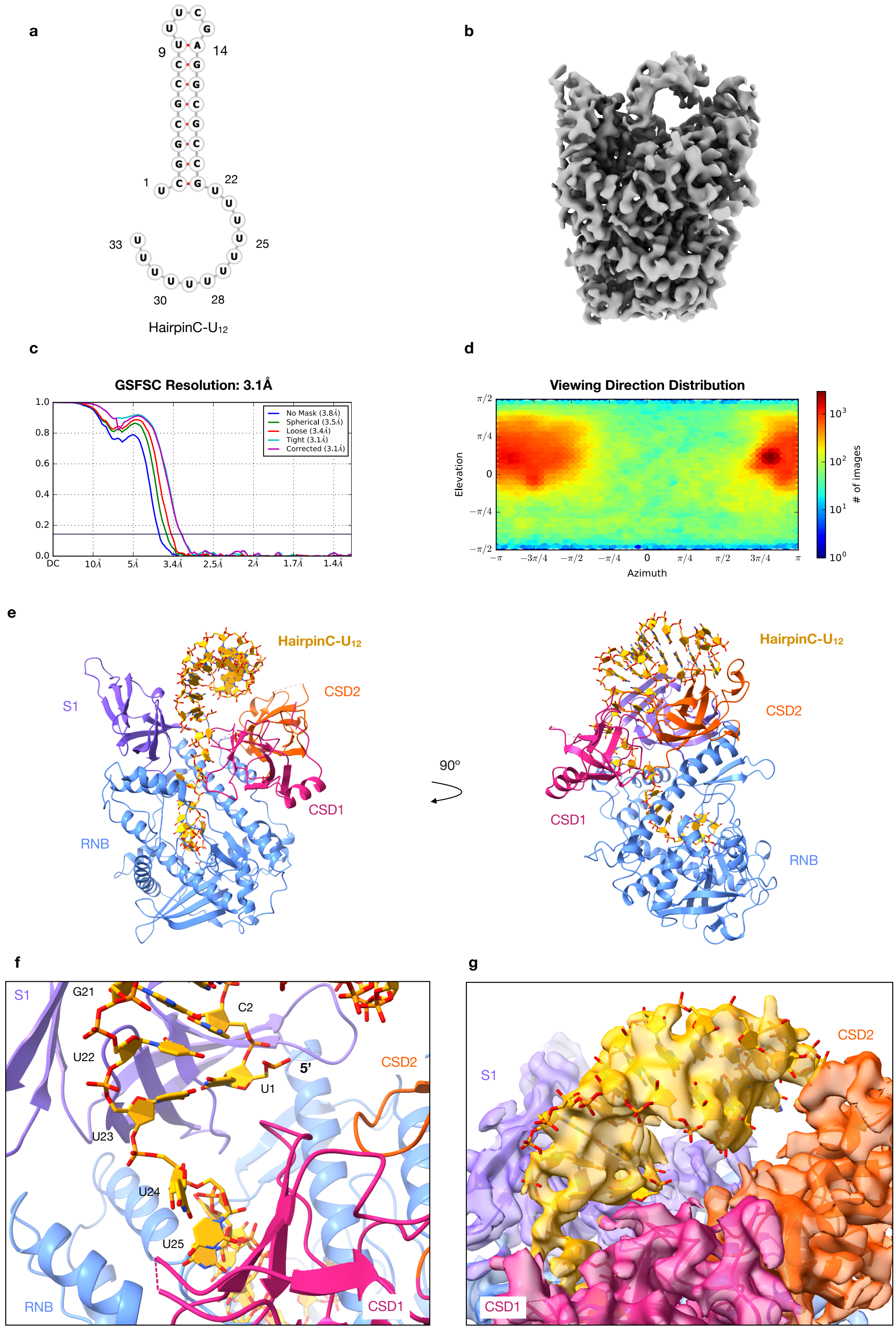
a
9
14
1
22
33
25
30
28
HairpinC-U12
b
c
GSFSC Resolution: 3.1Å
No Mask (3.8Å)
Spherical (3.5Å)
Loose (3.4Å)
Tight (3.1Å)
Corrected (3.1Å)
DC
10Å
5Å
3.4Å
2.5Å
2Å
1.7Å
1.4Å
d
Viewing Direction Distribution
Elevation
Azimuth
### of images
e
HairpinC-U12
S1
CSD2
CSD1
RNB
90º
HairpinC-U12
CSD2
CSD1
RNB
f
S1
G21
C2
U22
5'
U1
CSD2
U23
U24
U25
RNB
CSD1
g
CSD2
S1
CSD1

**Extended data Fig. 3 Engaging double-stranded RNA by Dis3L2**. **a**, Predicted RNA-fold for hairpinC-$U_{12}$. **b**, Final 3D map of human Dis3L2 in complex with hairpinC-$U_{12}$ with **c,** FSC and **d,** the particle viewing direction distribution. The final 3D map was produced by homogeneous refinement of roughly 532,000 particles in cryoSPARC. **e,** Structure of Dis3L2 in complex with hairpinC-$U_{12}$, front and side view. **f**, Basal junction of stem and overhang. **g**, Overlay of map and structure showing position of double-helical stem of hairpinC-$U_{12}$.

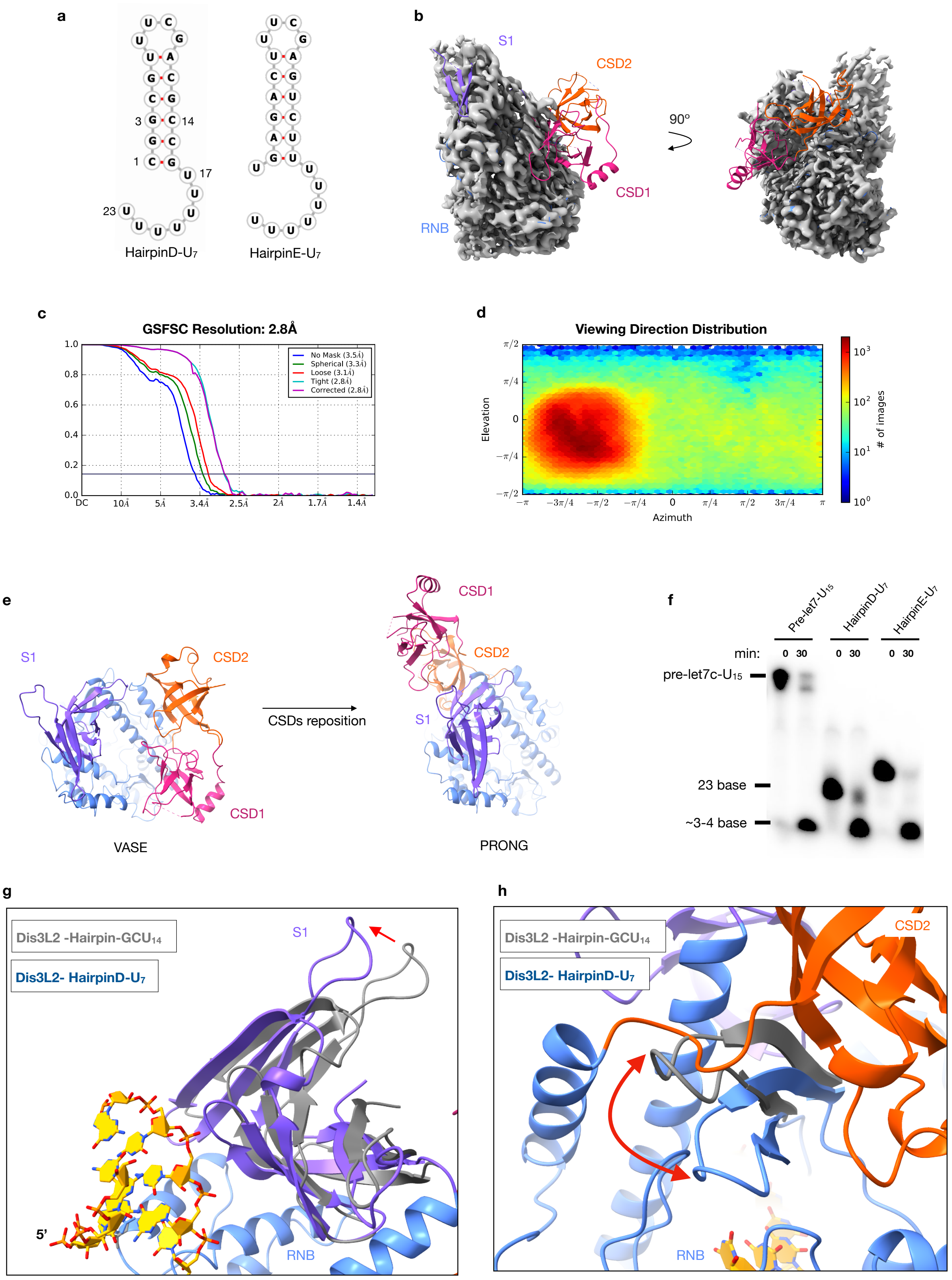
a
HairpinD-U7
HairpinE-U7
b
S1
CSD2
90°
CSD1
RNB
c
GSFSC Resolution: 2.8Å
No Mask (3.5Å)
Spherical (3.3Å)
Loose (3.1Å)
Tight (2.8Å)
Corrected (2.8Å)
d
Viewing Direction Distribution
Elevation
Azimuth
### of images
e
S1
CSD2
CSD1
CSDs reposition
VASE
PRONG
f
Pre-let7-U15
HairpinD-U7
HairpinE-U7
min:
0 30
pre-let7c-U15
23 base
~3-4 base
g
Dis3L2 -Hairpin-GCU14
Dis3L2- HairpinD-U7
S1
5'
RNB
h
Dis3L2 -Hairpin-GCU14
Dis3L2- HairpinD-U7
CSD2
RNB

**Extended data Fig. 4 Conformational change is triggered by structured RNA substrates with shortened 3' overhangs.** **a**, Predicted RNA-fold for hairpinD-$U_7$ (75%- GC content of dsRNA) and hairpinE-$U_7$ (50% GC content of dsRNA). **b**, Final 3D map of human Dis3L2 in complex with hairpinD-U7 superimposed with the structure of RNA-free Dis3L2 fit in the map. **c**, GSFSC and **d**, particle viewing direction distribution. The final 3D map was produced by homogeneous refinement of roughly 539,000 particles in cryoSPARC. **e**, View from the top, showing the change in position of the S1 and CSD domains in the vase and prong conformation. **f,** Urea-PAGE of nuclease activity assay against pre-let7-$U_{15}$, hairpinD-$U_7$ and hairpinE-$U_7$. Species length is indicated on the left of the gel. **g**, Difference in positioning of the S1 domain and **h,** difference in positioning of a RNB hairpin (residues 555-572) in the alignment of hairpin-$GCU_{14}$ and hairpinD-$U_7$ Dis3L2 structures.

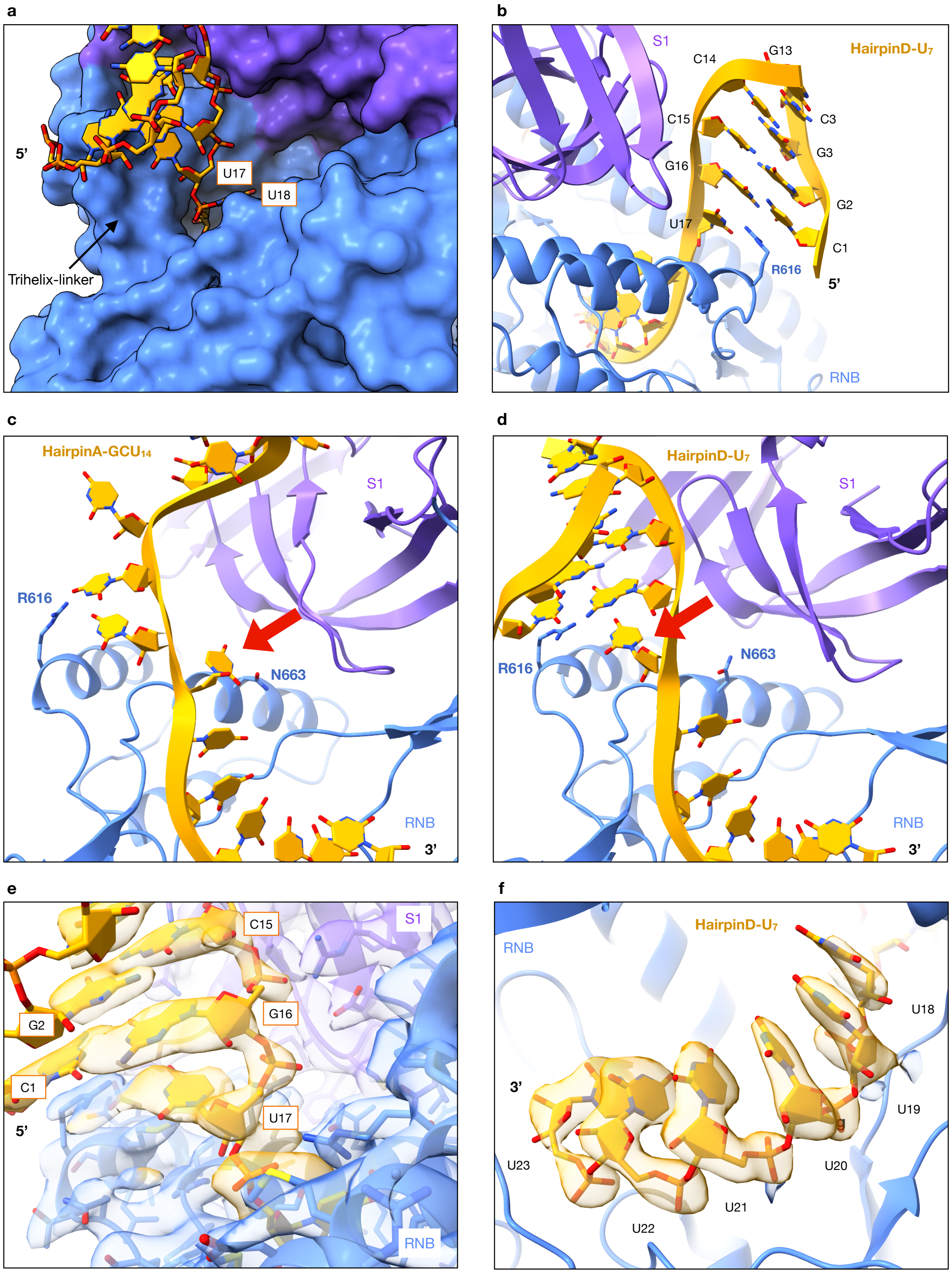

**Extended Data Fig. 5 The base of the double-helical stem is positioned above an RNB trihelix-linker. a,** The trihelix-linker forms the final barrier to the dsRNA before the tunnel to the active site. **b**, R616 on the junction of the trihelix stacks under C1 and points towards the first base of the 3'overhang (U17). **c**, The seventh base from the 3'end (red arrow) points towards N663 in the vase conformatio**n, d**, and flips towards R616 in the prong conformation. **e**, Overlay of the cryoEM map of the HsDis3L2-

hairpinD-$U_7$ structure at the hairpin basal junction. **f**, 5-nt of the $U_7$ overhang are buried in the tunnel of the RNB.

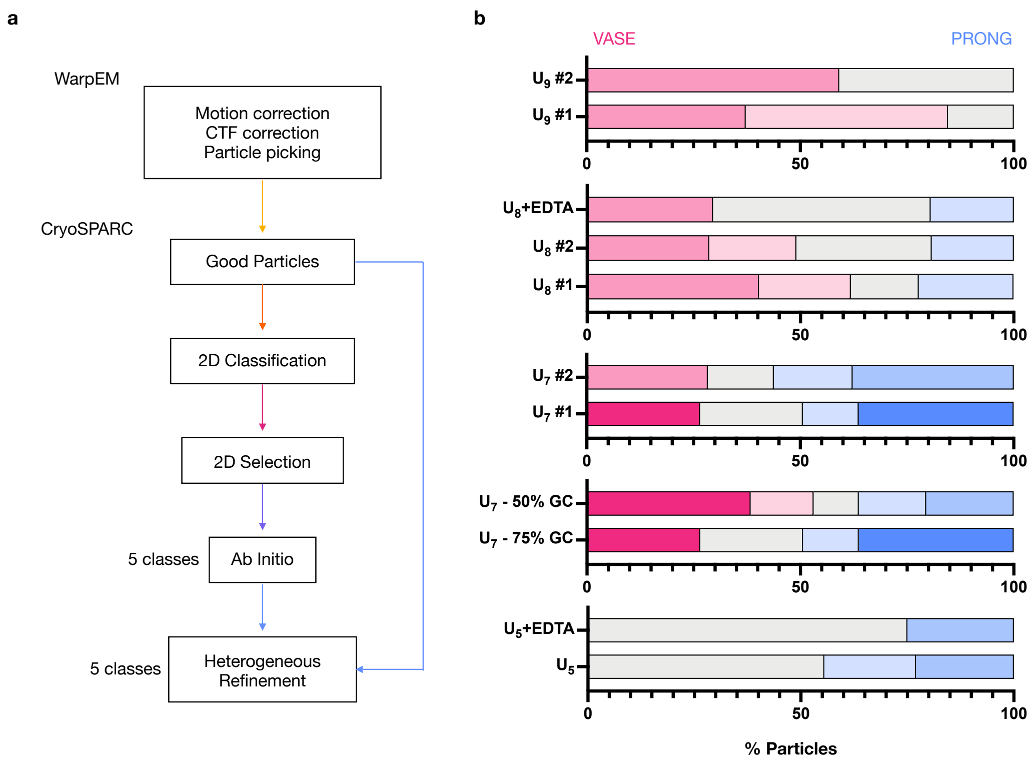

**Extended data Fig. 6 CryoEM 3D class distribution of particles. a**, Standardized data processing workflow to compare 3D class distribution of particles (Methods). Good particles picked by WarpEm's neural network-based picker first underwent 2D classification in cryoSPARC. Particles from classes with high estimated resolution and particle number were selected for *ab initio* reconstruction with 5 classes (Methods). The resulting 5 initial models were then used as starting references in a heterogeneous refinement with the full set of good particles. The resulting refinements were evaluated for their similarity to the prong or vase conformation and the proportion of particles belonging to each class plotted in **b**. **b**, Comparison of particle distribution in different datasets. Numbers #1 and #2 indicate repeat data collections of the same complex. Controls with EDTA (100 $\mu$M) were done for the hairpinD-$U_8$ and -$U_5$ complexes. Effect of hairpin GC content on conformational distribution of Dis3L2 was evaluated for a -$U_7$ overhang. X-axis: % of particles in vase (pink) or prong (blue) conformation, y-axis: individual datasets RNA-free or hairpin RNA-bound Dis3L2, numbers denote the length of the 3' overhang. The deeper color indicates higher quality 3D-reconstructions, gray indicates particles that did not contribute to a meaningful reconstruction.

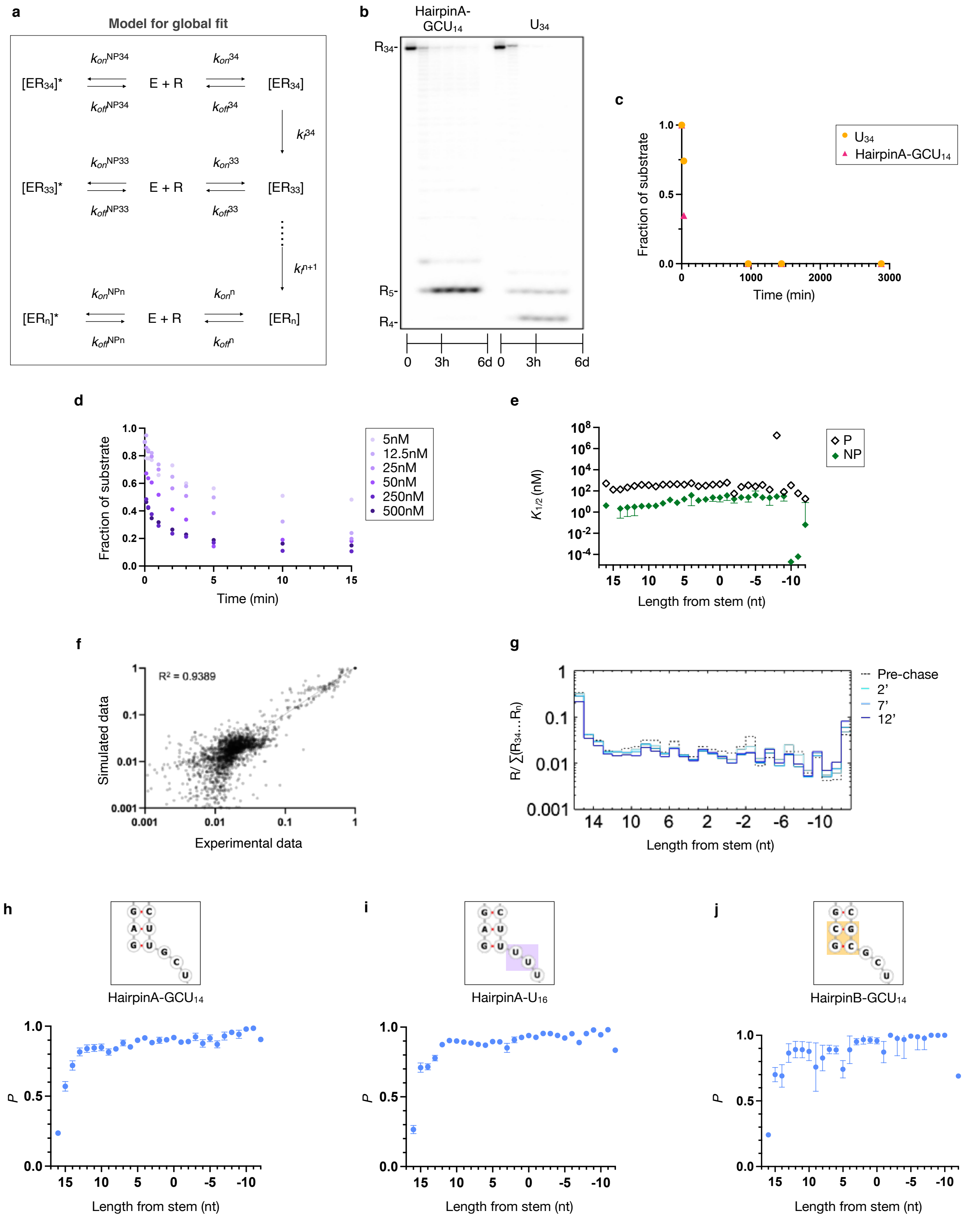
a
Model for global fit
[ER34]*
E + R
[ER34]
[ER33]*
E + R
[ER33]
[ERn]*
E + R
[ERn]
b
HairpinA-GCU14
U34
R34-
R5-
R4-
0
3h
6d
c
Fraction of substrate
Time (min)
U34
HairpinA-GCU14
d
Fraction of substrate
Time (min)
5nM
12.5nM
25nM
50nM
250nM
500nM
e
K1/2 (nM)
Length from stem (nt)
P
NP
f
R² = 0.9389
Simulated data
Experimental data
g
R/∑(R34...Rn)
Length from stem (nt)
Pre-chase
2'
7'
12'
h
HairpinA-GCU14
P
Length from stem (nt)
i
HairpinA-U16
P
Length from stem (nt)
j
HairpinB-GCU14
P
Length from stem (nt)

**Extended data Fig. 7 Kinetic analysis of WT HsDis3L2 against hairpin RNAs. a**, Minimal kinetic model used for global data fitting. **b**, Gel showing degradation of hairpinA-GCU$_{14}$ and ss-U$_{34}$ to completion by Dis3L2 over longer time course. **c**, Quantification of substrate disappearance from reaction shown in b. **d**, Fraction of hairpinA-GCU$_{14}$ substrate disappearance measured over time at various Dis3L2 concentrations. **e**, Comparison of functional equilibrium dissociation constants for productive ($K_{1/2}^{P}$) and non-productive ($K_{1/2}^{NP}$) binding determined in the global fit of WT Dis3L2 on hairpinA-GCU$_{14}$ (the data point for x=15 was omitted due to large uncertainty). **f**, Correlation of experimental data vs the corresponding data calculated using the kinetic parameters for pre-steady state reactions of Dis3L2 on hairpinA-GCU$_{14}$. **g**, Step-plot showing the change in substrate and intermediate species at discrete timepoints during pulse-chase reaction at 50nM Dis3L2 on hairpinA-GCU$_{14}$, the initial timepoint was taken before addition of chase at 3min. **h-j**, Comparison of Dis3L2 processivity on hairpinA-GCU$_{14}$, hairpinA-U$_{16}$, and hairpinB-GCU$_{14}$ ($P^{\text{HairpinB-GCU14}}$, x= -11 was omitted due to large uncertainty). Error bars are shown as vertical lines. $K_{1/2}$ errors (**e**) were calculated from SEM of $k_{off}$ and $k_{on}$. Processivity error for **i** represents the SEM, while **h** and **j** were calculated from the SEM of $k_f$ and $k_{off}$ (Methods),

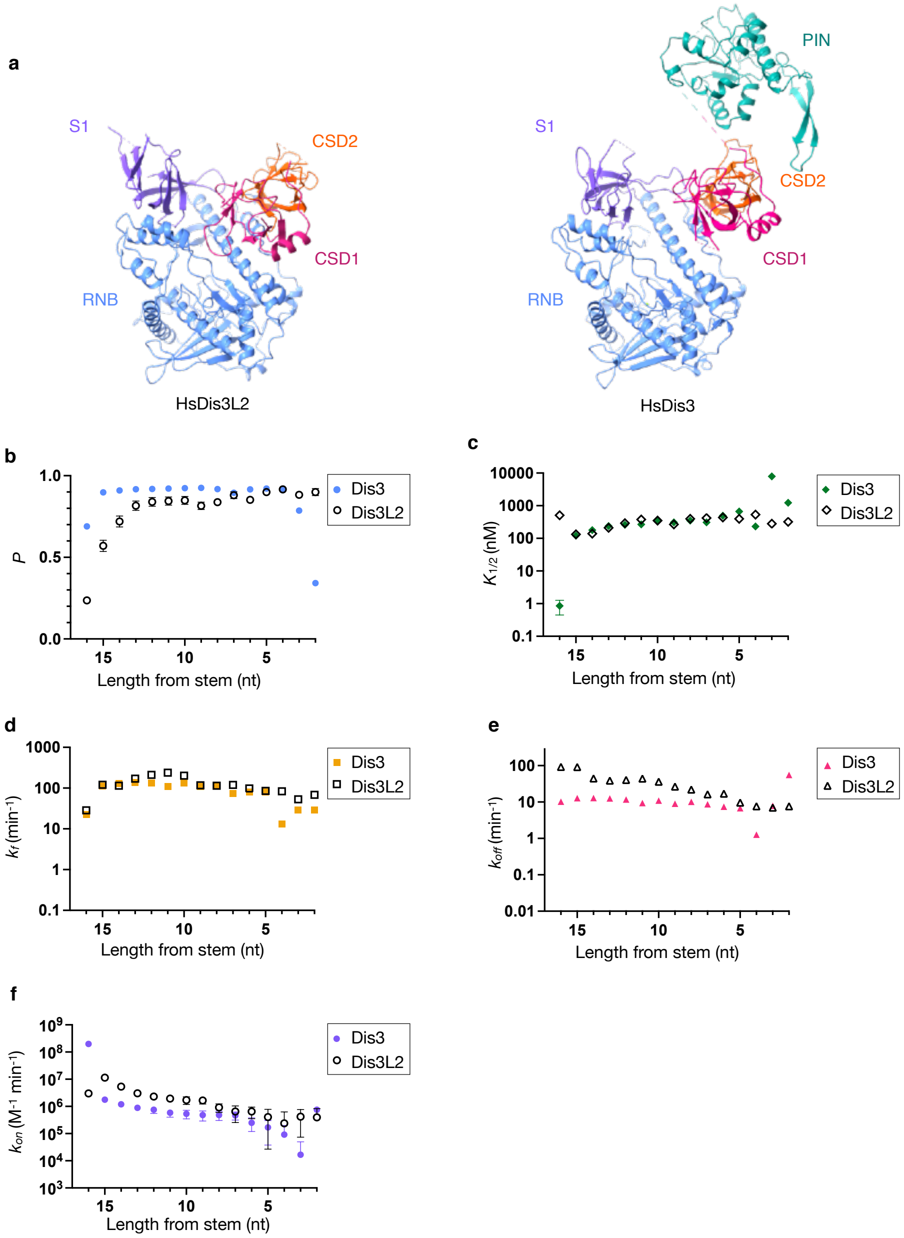

**Extended data Fig. 8 Comparison of structures and kinetic profiles of human Dis3 and Dis3L2. a**, Structures of Human Dis3L2 and Human Dis3 (PDB: 6d6q)[54]. Kinetic profile for hairpinA-GCU$_{14}$ degradation by HsDis3 at single-nt resolution **b**, Processivity (blue circles), **c**, Functional equilibrium dissociation constant for productive binding ($K_{1/2}$) (green diamonds), **d**, forward rate constant (orange squares), **e**, dissociation rate constant (pink triangles), and, **f**, association rate constants (purple circles) compared to HsDis3L2 (gray). Error bars are shown as vertical lines, and represent the SEM (**d**-**f**). Processivity errors for **b** were calculated from the SEM of $k_f$ and $k_{off}$, while $K_{1/2}$ errors (**e**) were calculated from SEM of $k_{off}$ and $k_{on}$ (Methods).

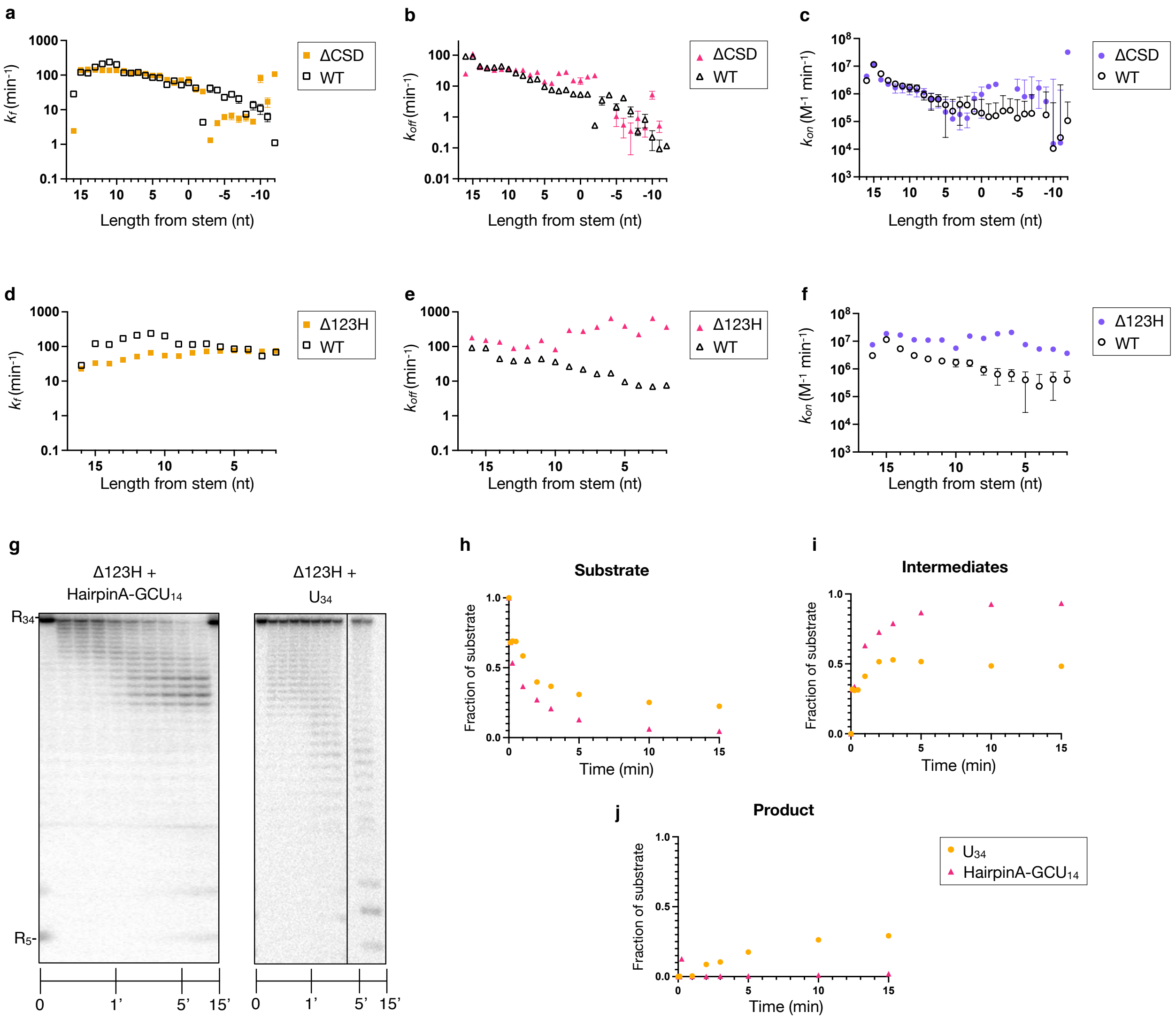

**Extended data Fig. 9 Contribution of the CSDs and trihelix-and-linker to degradation of structured substrates by human Dis3L2.** Kinetic profile for hairpinA-GC-$U_{14}$ degradation by ΔCSD mutant at single-nt resolution **a**, forward rate constants (orange squares), **b**, dissociation rate constants (pink triangles), and, **c**, association rate constants (purple circles) compared to WT Dis3L2 (gray), (the datapoints for $k_{on}$ at x=-3 and -4 were omitted due to large uncertainty). Kinetic profile for hairpinA-GC-U14 degradation by Δ123H (trihelix deletion) mutant at single-nt resolution **d**, forward rate constants (orange squares), **e**, dissociation rate constants (pink triangles), and, **f**, association rate constants (purple circles) compared to WT Dis3L2. **g**, Representative gels showing the degradation profile by 250 nM Δ123H mutant against 1 nM hairpinA-$GCU_{14}$ and $U_{34}$ substrates. **h-j**, Fraction of substrate disappearance, fraction of intermediates, and fraction of end-product accumulation over time from reactions shown in **g**. Error bars are shown as vertical lines, and represent the SEM (**a**-**f**).

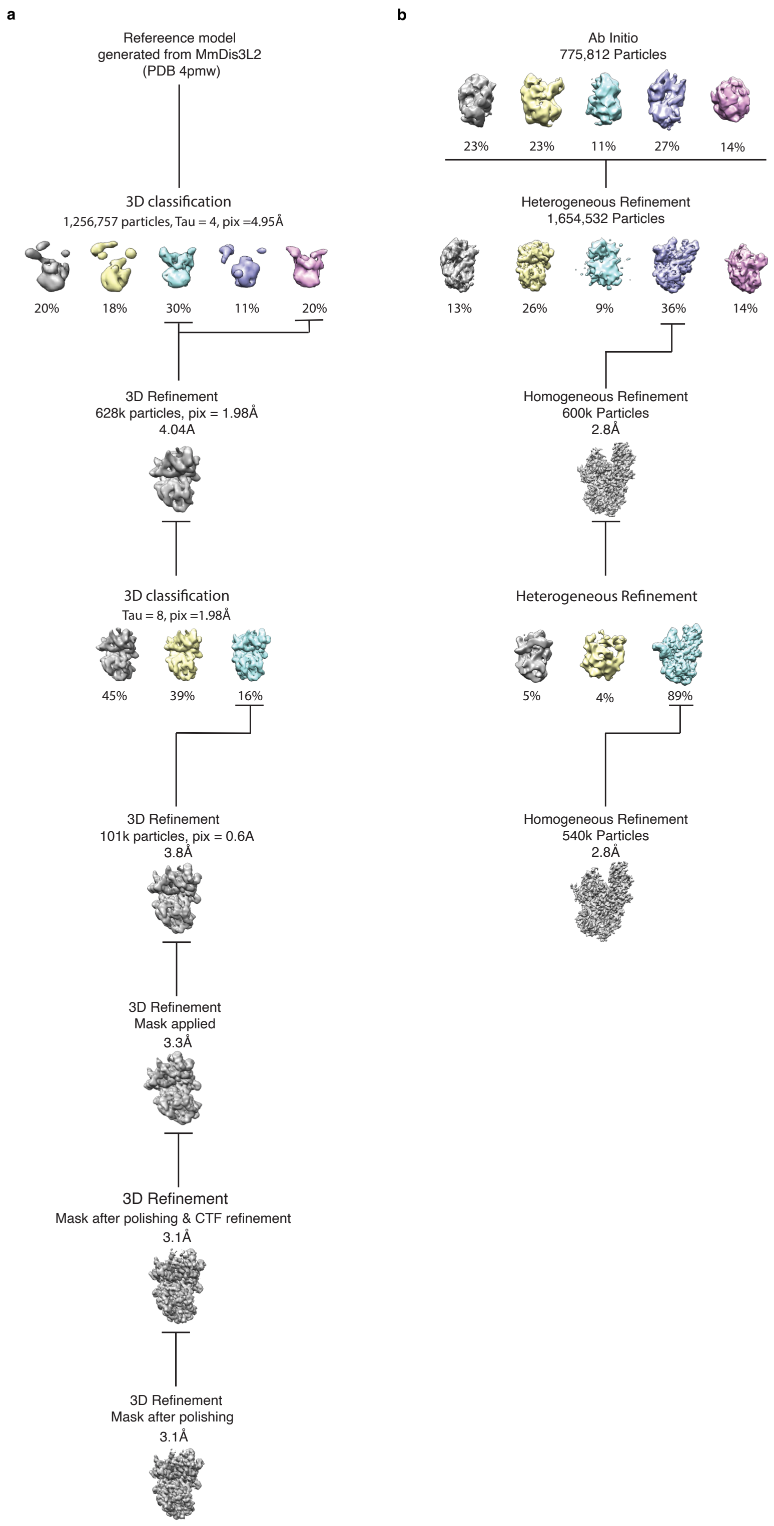

**Supplementary Figure 1. Representative cryoEM data processing pathways.** All datasets were pre-processed in WarpEM including motion correction, CTF estimation and particle picking. Further processing was done in Relion or CryoSPARC. **a**, Data processing of HsDis3L2-hairpinA-GCU$_{14}$ dataset in Relion. Following 2D classification to remove junk/bad particles the structure of MmDis3L2 was used to generate a starting model, and filtered to 20Å. A series of refinement steps were then done to arrive at the final 3.1Å resolution model used to build the HsDis3L2-hairpinA-GCU$_{14}$ structure. **b**, Data processing of the HsDis3L2-hairpinD-U$_7$ dataset was done in CryoSPARC.

Following 2D classification, a subset of the best particles (as determined by high resolution estimation and high particle number in the 2D classes) were used for *ab initio* reconstruction of 5 classes. The resulting models were then used as starting references in a heterogeneous refinement using all the good particles picked by WarpEM. The highest resolution class was further refined in hetero- and homogeneous refinements to a final resolution of 2.8Å.

**Supplementary Table 1. Cryo-EM data collection, refinement and validation statistics**

| | RNA-free HsDis3L2$^{D391N}$ | HsDis3L2 with hairpinA-GCU$_{14}$ | HsDis3L2 with hairpinC-U$_{12}$ | HsDis3L2 with hairpinD-U$_7$ |
|---|---|---|---|---|
| **Data collection and processing** | | | | |
| Magnification | 215,000X | 215,000X | 130,000X | 130,000X |
| Voltage (kV) | 300 | 300 | 300 | 300 |
| Electron exposure (e–/Å$^2$) | 2.07 | 2.07 | 2.08 | 2.08 |
| Defocus range (µm) | -2.8 to -0.5 | -2.8 to -0.5 | -2.8 to -0.5 | -2.8 to -0.5 |
| Pixel size (Å) | 0.84 Å | 0.84 Å | 0.64 Å | 0.64 Å |
| Symmetry imposed | / | / | / | / |
| Initial particle images (no.) | 990,287 | 1,256,757 | 1,375,384 | 1,654,532 |
| Final particle images (no.) | 162,793 | 101,028 | 531,561 | 539,985 |
| Map resolution (Å) 0.143 FSC threshold | 3.4 | 3.1 | 3.1 | 2.8 |
| **Refinement** | | | | |
| Refinement program and final refinement | CryoSPARC Homogeneous refinement | RELION Refine 3D | CryoSPARC Non-uniform refinement | CryoSPARC Homogeneous refinement |
| FSC (model) 0/0.143/0.5 | | | | |
| Masked | 3.3/3.4/3.6 | 3.0/3.1/3.2 | 3.0/3.2/3.4 | 2.7/2.8/2.9 |
| Unmasked | 3.3/3.4/3.8 | 3.1/3.1/3.3 | 3.1/3.3/3.5 | 2.8/2.8/3.0 |
| Map sharpening *B* factor (Å$^2$) | -150.7 | -79 | -160.4 | -109.8 |
| Model composition | | | | |
| Atoms | 5371(H: 0) | 5789 (H: 0) | 6197 (H: 0) | 5768(H: 0) |
| Residues | Protein: 676 Nucleotide: 0 | Protein: 690 Nucleotide: 15 | Protein: 692 Nucleotide: 33 | Protein: 686 Nucleotide: 15 |
| *B* factors (Å$^2$) | | | | |
| Protein | 41.65/137.90/86.19 | 16.07/113.65/45.17 | 37.03/150.05/75.79 | 39.39/141.89/71.65 |
| Nucleotide | / | 38.44/86.94/55.20 | 60.52/215.31/163.96 | 51.96/143.50/104.02 |
| R.m.s. deviations | | | | |
| Bond lengths (Å) | 0.007 (0) | 0.004 (0) | 0.011 (0) | 0.004 (0) |
| Bond angles (°) | 1.343 (2) | 0.837 (0) | 1.093 (0) | 0.699 (0) |
| Validation | | | | |
| MolProbity score | 1.45 | 1.16 | 1.31 | 1.44 |
| Clash score | 7.14 | 3.75 | 4.46 | 6.65 |
| Poor rotamers (%) | 0.34 | 0 | 1.31 | 0 |
| Ramachandran plot | | | | |
| Favored (%) | 0 | 0 | 0 | 0 |
| Allowed (%) | 2.25 | 1.46 | 1.46 | 2.35 |
| Disallowed (%) | 97.75 | 98.54 | 98.54 | 97.65 |

**Supplementary Table 2. Asymmetric error analysis for WT Dis3L2 on hairpinA-GCU$_{14}$**

| Parameter | Units | Best Fit | Upper Bound | Lower Bound |
|---|---|---|---|---|
| $k_{on}^{P}$ | $nM^{-1}\ min^{-1}$ | 0.180 | 0.200 | 0.159 |
| $k_{off}^{P}$ | $min^{-1}$ | 92.0 | 107 | 86 |
| $k_{forward}$ | $min^{-1}$ | 28.5 | 32.0 | 25.5 |
| $k_{off}^{NP}$ | $min^{-1}$ | 0.125 | 0.152 | 0.0770 |

**Supplementary Table 3. Asymmetric error analysis for ΔCSD on hairpinA-GCU$_{14}$**

| Parameter | Units | Best Fit | Upper Bound | Lower Bound |
|---|---|---|---|---|
| $k_{on}^{P}$ | $nM^{-1}\ min^{-1}$ | 0.258 | 0.296 | 0.225 |
| $k_{off}^{P}$ | $min^{-1}$ | 25.2 | 28.9 | 21.9 |
| $k_{forward}$ | $min^{-1}$ | 2.44 | 2.74 | 2.15 |
| $k_{off}^{NP}$ | $min^{-1}$ | 0.0805 | 0.129 | 0.0338 |

**Supplementary Table 4. Asymmetric error analysis for Δ123H on hairpinA-GCU$_{14}$**

| Parameter | Units | Best Fit | Upper Bound | Lower Bound |
|---|---|---|---|---|
| $k_{on}^{P}$ | $nM^{-1}\ min^{-1}$ | 0.446 | 0.477 | 0.423 |
| $k_{off}^{P}$ | $min^{-1}$ | 181 | 196 | 169 |
| $k_{forward}$ | $min^{-1}$ | 22.8 | 25.0 | 21.9 |
| $k_{off}^{NP}$ | $min^{-1}$ | 0.281 | 0.331 | 0.19 |

#### Supplementary Discussion

##### A disordered CSD1 loop insertion with possible regulatory roles

Dis3L2 possesses a roughly 120 amino acid long loop in CSD1 (L115-R231) that has not been resolved in any Dis3L2 structures to date. Interestingly, the pseudo-nuclease Ssd1 and ScDis3 also have a long loop insertion in CSD1[49,50]. In both Ssd1 and ScDis3, part of this loop forms a helix. In Ssd1 the helix sits in-between the CSDs and S1 domain, blocking the apical face of the protein. In the case of ScDis3, the CSD1 insertion has been observed in two different positions depending on the way Dis3 associates to the rest of the RNA exosome. In one conformation, referred to as 'open', this helix lies in between the S1 and CSDs covering the apical face like a lid[50], while in the second, or 'closed' conformation, the CSD1 loop and helix reposition to expose the apical face, accompanied by a widening of the CSDs and S1 domains (HsDis3: 6h25, 6d6q). In contrast, in human Dis3 and bacterial RNase R and RNase II this region is much shorter, largely unresolved and has little sequence conservation (RNase R: 5xgu, RNase II: 2ix1). Examination of our RNA-free HsDis3L2 EM maps revealed undefined density between the CSDs and S1 domains, which suggests that the CSD1 loop region is flexible and interacts transiently with the apical face.

While the significance of this loop insertion in Dis3L2 is unknown, it is possible that it plays a functional role, for example by modulating RNA access and activity or influencing interactions with other factors. Furthermore, proteomics studies have previously identified phosphorylation sites in the CSD1 loop region[51]. These sites could play a role in the regulation of RNA binding, cellular localization and/or recognition of other factors such as the TUT enzymes.

**Non-productive binding and non-nuclease roles of Dis3L2**

Non-productive binding describes binding events in which the association between the protein and RNA is not conducive to degradation: the RNA does not bind fully into the active site, the RNA 3' end is misoriented or the RNA is bound but the protein is in a non-productive conformation.

An example of non-productive binding was observed in the crystal structure of MmDis3L2. While the RNA substrate provided was only 13 nucleotides in length, 14 nucleotides of density were seen due to two different positions of the RNA: either with the 3' end either right in the active site or removed away by one nucleotide, leaving the active site open. The latter position represents a non-productive state. Non-productive binding has been measured in other nucleases such as RRP6[48].

Our global kinetic model identified a contribution from non-productive binding during substrate processing by HsDis3L2. Surprisingly, in the initial binding step, non-productive binding is 10-fold tighter compared to productive binding (Extended data Fig. 7e). Furthermore, our analysis has shown that the CSDs provide roughly half of the non-productive binding affinity (non-productive $K_{1/2}$ of 4.2 $\pm$ 0.41 (WT) vs. 8.5 $\pm$ 0.73 ($\Delta$CSD) nM) and removing them improves productive binding five-fold (productive $K_{1/2}$ of 508.3 $\pm$ 1.27 (WT) vs. 97.7 $\pm$ 0.35 ($\Delta$CSD) nM). This suggest that along with mediating association with substrates targeted for degradation, Dis3L2's RNA-binding regions may play a role in non-nucleolytic RNA-binding functions. To confirm that the non-productively bound species are not due to non-cleavable RNA, we performed long time-course experiments and showed that all species do degrade to the 5 to 4 nt end product (Extended data Fig. 7b-c).

The large contribution of non-productive binding might indicate that Dis3L2 functions in other roles which do not require nuclease activity. A recent study of hepatocellular carcinoma (HCC) appears to have identified one such case[8]. Dis3L2 was found to be highly expressed in HCC tissues and promoted alternative splicing of the Rac1 gene through a nuclease-independent mechanism. Dis3L2 was shown to bind the Rac1 pre-mRNA via the S1 domain and recruit hnRNP-U through its CSDs. This enabled the production of Rac1b, an isoform that promotes transformation and tumorigenesis[8].

Evolutionary analysis of Dis3L2 has revealed that the protein has lost nuclease activity at least four times during fungal evolution, while the CSDs have remained conserved, thereby suggesting a role for the protein outside of RNA degradation[52]. The Dis3L2 homolog in *S. cerevisiae*, Ssd1, is an example of this, losing both canonical RNase II/R catalytic residues and acquiring a loop insertion that blocks the tunnel to the active site. Ssd1 has been reported to act as a translational repressor of certain mRNAs involved in cell growth and cytokinesis, and deletion of Ssd1 was found to have pleiotropic effects on stress tolerance[52,53].

**Role of the CSDs in degradation of structured RNA substrates**

Analysis of a Dis3L2 mutant in which the CSDs had been removed, ΔCSD, showed an impact on both the first catalytic step as well as the initial unwinding phase (Extended data Fig. 9a, b). Moreover, prior to duplex unwinding, the dissociation constant increased (Extended data Fig. 9b). Our structural data shows that the CSDs switch to the prong conformation at overhang lengths of roughly 8 nt (Fig. 1e, g), which makes the fact that we see an impact on binding during the unwinding phase in the ΔCSD somewhat confusing. There are two models that might explain these observations. The CSDs could be contributing to binding indirectly, by stabilizing the repositioning of the S1 domain to directly interact with the RNA double-helix in the prong conformation. Alternatively, the CSDs may swing back to bind the dsRNA directly. Since cryoEM analysis showed that a hairpinD-5U substrate led to the prong conformation alone, the former model seems more likely. However, active turnover conditions may allow for more dynamic back-and-forth movement of the CSDs during degradation.

**dsRNA unwinding by Dis3L2**

Our structural data suggests that strand unwinding has to take place around a 3' overhang length of roughly 5 nt – as this is the number of nucleotides buried in the narrow tunnel to the active site in the prong conformation. Analysis of WT Dis3L2 degrading hairpinA-GC14U revealed that after the first nucleotide cleavage step, the processivity does not vary significantly leading up to the start of the double-helix. Though small dips in the processivity can be observed (for example at 9-, 6- and 3 nt overhang) that might correlate with the enzyme's close encounter with the dsRNA, it is difficult to attribute these to specific events. Since there is a weak base-pair (GU and AU) at the base of the hairpinA stem, we designed a substrate in which this was changed to GC and CG respectively (Extended data Fig. 7j, hairpinB-GC14U). We analyzed the kinetic profile with hairpinB-GC14U and found that while the overall pattern was largely the same, some key points displayed more pronounced changes. Most interestingly, we see a significant dip in the processivity at an overhang length of 5 and 9 nt (though the error for the latter is larger). These may reflect some stalling prior to the protein's large conformational change (9 to 8 nucleotides) and dsRNA unwinding (5 to 4 nucleotides).

#### RNA unwinding in related RNase II/R nucleases

*S. cerevisiae* has only one active Dis3 homolog which functions both in the cytoplasm and the nucleus and, in contrast to its human Dis3 homolog, is able to independently degrade dsRNA. As mentioned earlier, ScDis3 has been observed to bind the exosome in two different conformations: open and closed. These conformational changes are distinct from what we have observed with Dis3L2. In the open conformation of ScDis3, single-stranded RNA bypasses the Exo9 core and enters directly into the nuclease. ScDis3 is positioned on the exosome core such that the RNA entry channel is solvent accessible and, as its apical face is obstructed by the CSD1 loop insertion, the RNA enters laterally from below the CSDs[50]. DsRNA entering this way might encounter the trihelix-linker more directly. However, if we assume that duplex unwinding requires direct interaction of the RNA hairpin-stem base with the trihelix-linker (as appears to be the case for Dis3L2), the CSDs would most likely still have to move, since their current position provides an opening of only ~13Å. Interestingly, in the closed conformation RNA passes through the exosome core channel to get to ScDis3 and the nuclease is rotated to position the lateral entry channel closer to the exosome core[50]. This conformational change is accompanied by a widening of the S1 and CSDs and a repositioning of the CSD1 loop insertion such that the apical face of the nuclease is now sterically accessible, similar to the structures of HsDis3 and Dis3L2[6,54,55]. In this conformation, alignments suggest that dsRNA could be accommodated on the apical face. However, to position the RNA over the trihelix-linker, as seen in the hairpin-7U structure, the CSDs would still have to move to avoid a steric clash. Single-molecule experiments measuring strand separation of a GC rich dsRNA by ScDis3 found that unwinding took place in 4 nucleotide bursts using the buildup of elastic energy from multiple backbone hydrolysis events [56]. In this model, it is suggested that CSD1 might stabilize the duplex prior to unwinding, at which point it would move out of the way.

The most compelling evidence for a conserved unwinding mechanism among other nucleases in the RNase II/R family, is perhaps the finding that RNase R also requires the trihelix-linker to degrade structured RNA substrates [30]. Deletion of the trihelix-linker region in RNase R resulted in a similar build-up of dsRNA intermediate products as we have observed in our nuclease assays of Δ123H. The structures of RNase R have shown that it also adopts a vase conformation. As no substrate-bound structures exist, it is currently unclear whether the RNA enters apically or laterally. Alignments of the structures of Dis3L2- hairpin-7U and RNase R show that the duplex of the RNA clashes with CSD1, suggesting that if a similar mechanism is employed, RNase R would also have to undergo a conformational change. In this regard, evidence for a conserved 'mobility' of the CSDs, at least amongst the Dis3L2 nucleases, comes from the structure of *S. pombe* Dis3L2 (SpDis3L2). In this RNA-free structure the CSDs are swung-out relative to the vase conformation, but only CSD2 could be resolved [57].
